# Profiling drug sensitivity of leukemic stem cells via bulk-to-single-cell deconvolution

**DOI:** 10.1101/2023.05.10.540140

**Authors:** Alexandre Coudray, Romain Forey, Benjamin Bejar Haro, Filipe Martins, Joana Carlevaro-Fita, Shaoline Sheppard, Sandra Eloise Offner, Gioele La Manno, Guillaume Obozinski, Didier Trono

## Abstract

*Ex-vivo* drug sensitivity screening allows the prediction of cancer treatment effectiveness in a personalized fashion. However, it only provides a readout on mixtures of cells, potentially occulting important information on clinically relevant cell subtypes. To address this shortcoming, we developed a machinelearning framework to decompose drug sensitivity recorded at the bulk level into cell subtype-specific drug sensitivity. We first determined that our method could decipher the cellular composition of bulk samples with top-ranking accuracy across five cancer types compared to state-of-the-art bulk deconvolution methods. We emphasize its effectiveness in the realm of Acute Myeloid Leukemia, where it appears to offer the most precise estimation of leukemic stem cell fractions across three test datasets and three patient cohorts. We then optimized an algorithm capable of estimating cell subtype- and single-cell-specific drug sensitivity, which we evaluated by performing *in-vitro* drug studies and in-depth simulations. We then applied our deconvolution strategy to the beatAML cohort dataset, currently the most extensive database of *ex-vivo* drug screening data. We developed a drug sensitivity profile tailored to specific cell subtypes, focusing on four therapeutic compounds predicted to target leukemic stem cells: the previously known midostaurin and A-674563, as well as SNS-032 and foretinib, which have not been previously linked to leukemic stem cells. Our work provides an attractive new computational tool for drug development and precision medicine.

## INTRODUCTION

*Ex-vivo* drug sensitivity screening (DSS) allows for the parallel testing of hundreds of compounds on cells extracted from a human tumor, potentially paving the way to personalized therapeutic decisions [1–3]. However, as currently performed, DSS assesses the susceptibility of cell mixtures and not of individual cell subtypes, hence fails to provide information that might be of utmost clinical importance, for instance on the drug sensitivity of rare cancer cells with stem-like properties. Coupling *ex-vivo* DSS with cell purification techniques such as fluorescence-activated cell sorting (FACS) can partly address this shortcoming, but sorting requires large quantities of cells, is limited by the availability of specific cell surface markers, and the procedure itself can impair the viability of some cell subtypes [4]. Therefore, computational methods capable of deconvoluting *ex-vivo* DSS would constitute attractive alternatives, as they would alleviate the need for cell subtypes purification.

Acute Myeloid Leukemia (AML) is a cancer type typically displaying extensive cellular heterogeneity, with the bone marrow of AML patients harboring a mixture of more than twenty cell subtypes, along with approximately 60 recurrently mutated genes and numerous rare mutations [5–7]. Conventional chemotherapies, involving anthracyclines and nucleoside analogs followed by bone marrow transplant, yield limited durable remission (around 20%) [8]. A persistent challenge in AML is drug resistance and relapse, urging a deeper grasp of the biological factors underpinning drug response. Notably, drug response frequently aligns with tumor cell maturation state, with certain drugs effective against less differentiated states [9–15], and others against more differentiated states [9, 12, 13, 16–23]. Recent single-cell sequencing further delineated AML cancer cell subtypes tied to differentiation state, including hematopoietic-stem-cell-like, progenitor-like, GMP-like, promonocyte-like, monocyte-like, and cDC-like types [24], several of which are directly relevant to the oncogenic process. Notably, the hematopoietic-like cancer cell subtype identified by Van Galen et al. (HSC-like) likely encompasses previously characterized leukemic stem cells (LSCs), thought to be responsible for relapse and decreased survival in AML patients [25–28]. Gene expression signatures from these leukemic stem cells (LSCs) also offer valuable prognostic insights [29–31], underscoring their clinical importance. Given this context, we posit that innovative approaches to elucidate the drug sensitivity of distinct cancer cell subpopulations could significantly enhance AML treatment strategies and potentially offer new avenues for targeting LSCs.

In this work, we introduce novel computational techniques that enable the concurrent estimation of cell subtype proportions and the deconvolution of *ex-vivo* drug sensitivity screening data into cell subtype- or single-cell-specific drug sensitivity profiles. We first developed CLIMB, a tool designed to decipher the cell subtype composition and expression within AML bulk samples, using a novel bulk-to-single-cell deconvolution approach. The outcome of CLIMB’s deconvolution serves as direct input for our subsequent method, CLIFF. CLIFF employs a probabilistic model to uncover latent variables corresponding to cell subtype survival probability, and thus deconvolute bulk-level drug sensitivity into cell-subtype-specific drug sensitivity profiles. We benchmarked our methods through the analysis of cancer cell line mixes we generated *in vitro*, and through pseudo-bulk deconvolution based on ten single-cell datasets across five cancer types. We then deployed CLIMB on three cohorts of AML patients, and used CLIFF to deconvolute cell-subtype drug sensitivity based on the beatAML cohort [7]. This led us to predict that the drugs A-674563, midostaurin, SNS-032, and foretinib would be effective in targeting HSC-like cancer cells. Further investigation of single-cell drug sensitivity predictions for venetoclax, a clinically used drug for AML, highlighted a subpopulation of venetoclax-resistant HSC-like cells.

## RESULTS

### Experimental design

Studying drug sensitivity in cancer cell subpopulations is essential yet challenging, employing various experimental approaches. Patient-derived xenograft models mimic patient heterogeneity and drug responses, but establishing them is resource-intensive. In contrast, *ex-vivo* drug sensitivity screening (DSS) assesses drug response in surgically extracted cancer cells, preserving their inherent diversity. Combined with fluorescence-activated cell sorting (FACS), DSS allows subpopulation-level analysis. However, FACS requires expensive antibodies and has limitations in isolating cell subtypes. Previous work by Bottomly et al. highlighted AML cell differentiation’s impact on drug sensitivity [32], while Zeng et al. linked HSC-like cell proportions (predicted using CiberSortX deconvolution method) to drug responses and outcomes [33]. In contrast, Karakaslar et al. showed that Monocyte-like content correlates with venetoclax resistance [34]. Yet, these studies relied solely on correlations between bulk-level drug sensitivity and cell subtype proportions from deconvolution methods. To enhance drug sensitivity deconvolution at the cellular level, we introduce a novel approach that decomposes bulk-level cell viability data into interpretable cell viability metrics at both cell subtype and single-cell resolutions.

Decoding drug sensitivity in cell subpopulations from heterogeneous mixtures requires knowledge or prediction of cellular composition. To address this challenge, we present CLIMB, a novel method that dissects the cellular composition of bulk samples (Fig. 1, upper part, and SFig. 1A, B). CLIMB employs a unique model that directly maps single cells from a reference dataset to the target bulk expression, aiming to optimize single-cell combinations. This approach makes CLIMB a signature matrix-free deconvolution method, sequentially mapping bulk samples to single cells and further to cell subtypes. Utilizing an iterative deconvolution process with scRNA-seq reference data and bulk RNA-seq expression as input, CLIMB can predict cell-subtype proportion and expression from a bulk RNA-seq sample. We utilized CLIMB to address a pivotal and clinically significant question: Can we accurately evaluate the fraction of leukemic stem cells contained in bulk samples from AML patients?

**Fig. 1.**
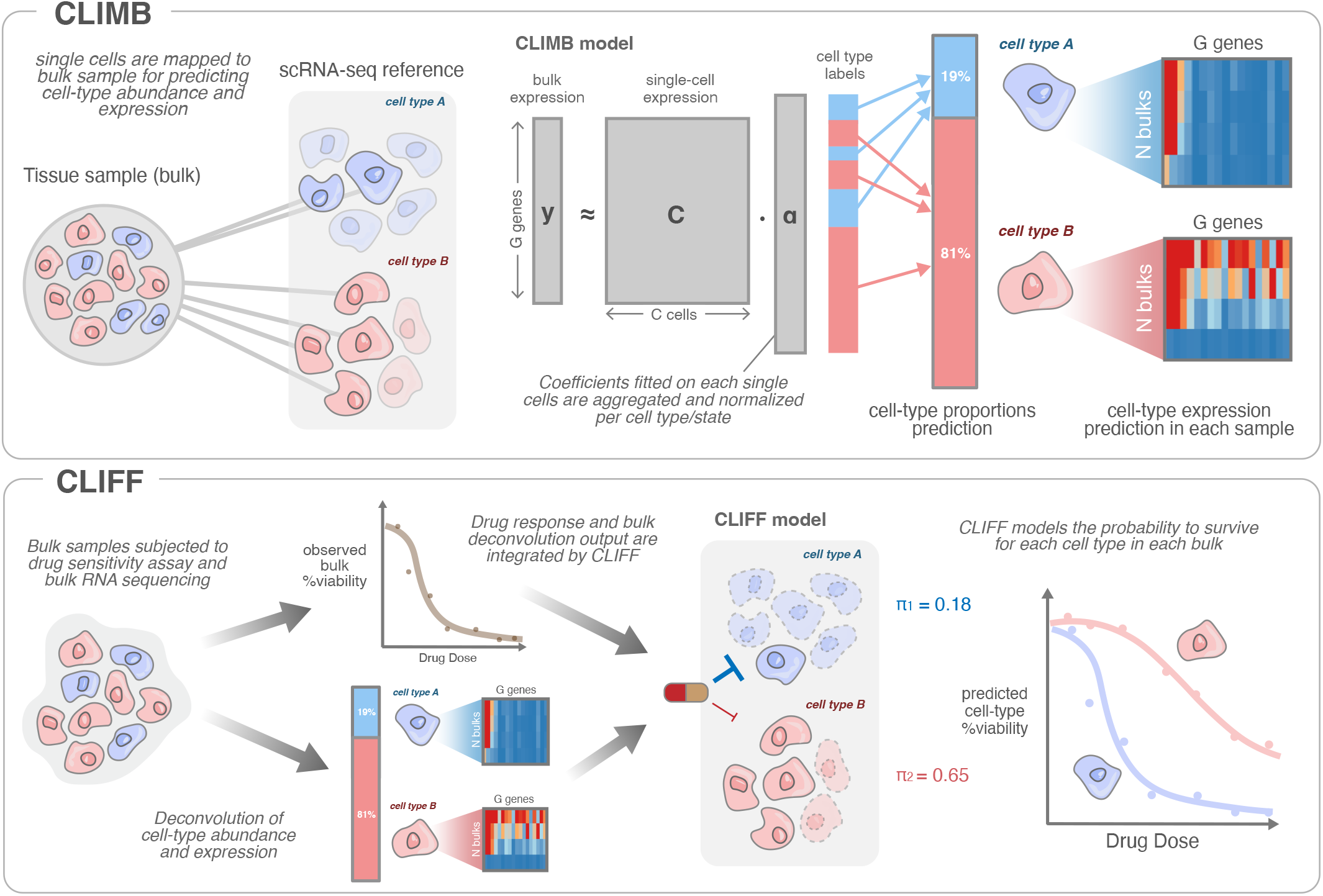
Experimental Design. (**Top**) CLIMB conducts bulk deconvolution using the full expression matrix from scRNA-seq, fitting coefficients on individual single cells, which are then aggregated at the cell-subtype level, allowing the inference of cell-subtype proportions and expression. (**Bottom**) Presented is the two-step process of our framework ‘CLIMB the CLIFF’. Initially, CLIMB deconvolution method is used to estimate cell subtype proportions and expression in each mixed sample. Subsequently, deconvoluted cell subtype abundance and expression are combined with drug sensitivity data to predict cell subtype drug sensitivity. In a nutshell, CLIFF employs a probabilistic model to uncover latent variables corresponding to cell subtype survival probability.

We further developed CLIFF, a probabilistic model that unveils latent variables corresponding to cell subtype cell viability, by leveraging cell-subtype proportions and expressions as input (Fig. 1, bottom, SFig. 1A). CLIFF’s framework defines cell viability as the ratio of surviving cells, adopting a Bernoulli distribution assumption for cell fate. CLIFF can thus directly integrate CLIMB’s output with cell viability assay data performed at the bulk level. Optionally, CLIFF incorporates somatic mutations as covariates and can predict drug sensitivity in ‘high-resolution’ mode by deconvoluting cell-subtype drug sensitivity within each bulk sample (SFig. 1B). We then tested CLIFF’s capability to predict the drug sensitivity of leukemic stem cells, aiming to assess susceptibilities of this crucial AML-specific cell subtype.

We introduce a dual approach, sequentially employing CLIMB and CLIFF, to dissect the cellular composition of tumor samples and unravel drug sensitivity at the cell subtype level, utilizing only bulk RNA-seq data along with bulk drug sensitivity information. Employing *in vitro* mixtures of human leukemic cell lines (HL60, SUDHL4, K562, and THP1), we assessed our bulk deconvolution in a known cell subtype fraction context, where individual cell line drug sensitivities can be experimentally determined, providing ground truth. By applying bulk RNA-seq alongside bulk *ex-vivo* drug screening data from the beatAML cohort, we translated bulk-level drug sensitivity into cell subtype-specific profiles. Our study then focus in a detailed exploration of the drug sensitivity specific to the leukemic stem cells, a cell subtype of utmost clinical significance.

### Evaluating Bulk Deconvolution Methods for Predicting Cellular Composition in Cancer Cell Line Mixtures

CLIMB infers cell-subtype proportions and can thus predict the composition of a cell mix (Fig. 2A). We generated *in vitro* pools containing various ratios of HL60, SUDHL4, K562, and THP1 human leukemic cells, analyzed by standard bulk RNA sequencing. We could thus use these pre-determined ratios as ground truth to compare the predictive value of CLIMB (Fig. 2B, C) and a panel of seven bulk deconvolution methods: BayesPrism [35], CiberSortX [36], TAPE [37], Scaden [38], MuSiC [39], BisqueRNA [40], and NNLS [39], using random proportions as a negative control. We then performed single-cell RNA sequencing of a mixture of the same cell lines, and used it as reference for all deconvolution methods with the same cell and gene selection (employing genes with higher estimated biological variance than technical variance, see Methods). The overall ranking among four metrics (average over 4 ranks), placed CLIMB and Scaden as the best deconvolution methods (Fig. 2D), followed by BayesPrism. Notably, CLIMB reached a higher accuracy than other deconvolution methods based on the root mean square error (RMSE, Fig. 2E). Scaden reached similar PCC, SCC, and R-squared, than CLIMB (Fig. 2C, right panel). A wilcoxon test taking four metrics into account showed significant advantage of CLIMB over other deconvolution methods, except Scaden. In summary, CLIMB demonstrated top-ranking accuracy in deconvoluting bulk RNA-seq samples from *in-vitro* cell mixtures, matching the performance of the deep learning-based method Scaden and surpassing other deconvolution methods.

**Fig. 2.**
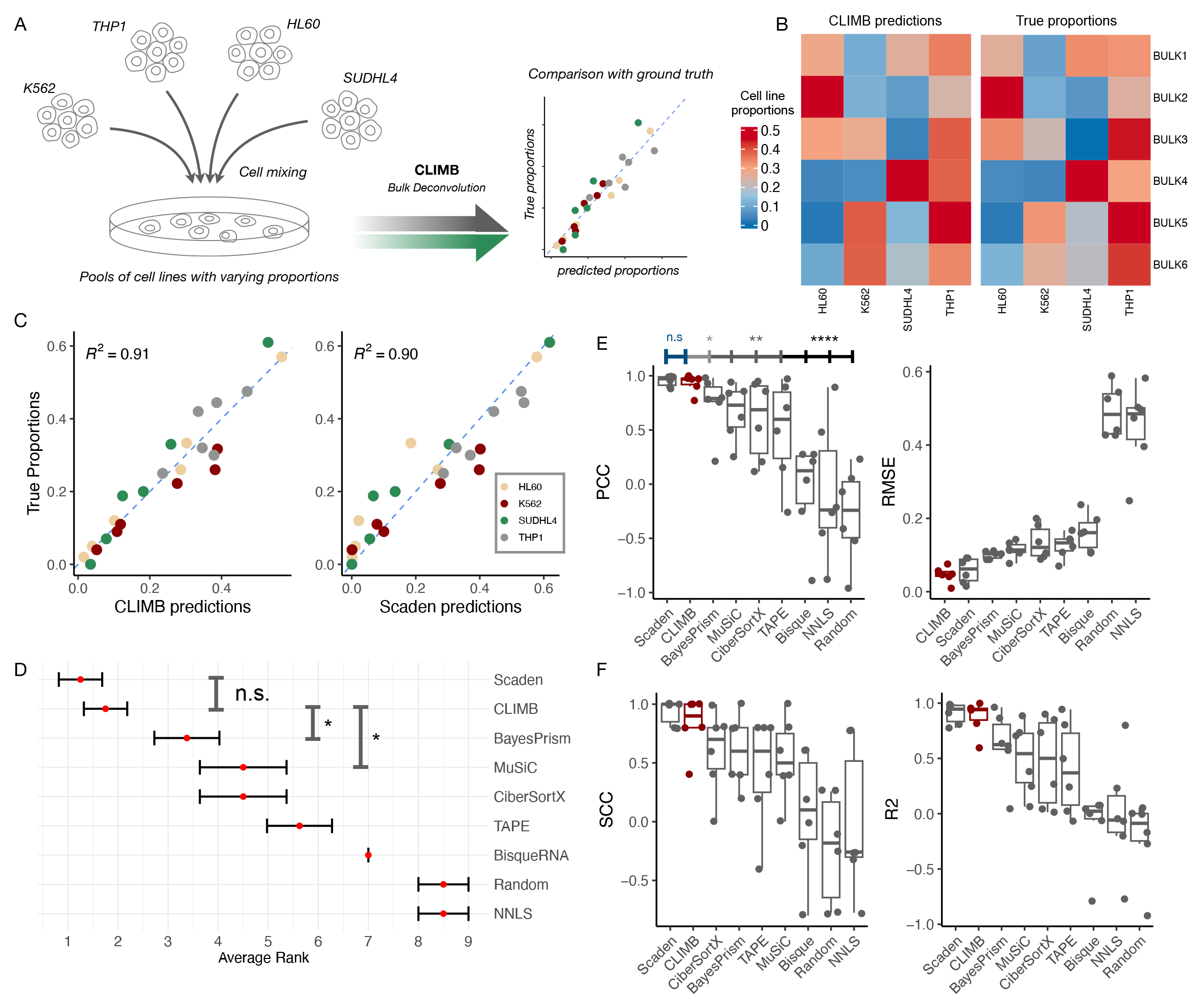
Cell Line Abundance Deconvolution using an *In-Vitro* Experiment with Known Proportions. (**A**) Experimental design for *in-vitro* experiment involving mixtures of human leukemic cell lines. (**B**) Heatmap displaying predicted cell line proportions by CLIMB (left) and actual proportions (ground truth, right). (**C**) Scatter plot comparing true versus predicted cell line abundances using CLIMB (left) and Scaden (right), with colors representing cell lines (6 bulk samples and 4 cell lines). (**D**) Overall results ranked based on median of four metrics (PCC, SCC, RMSE, and R-square) for bulk sample-level comparison. Wilcoxon rank test used for statistical comparison between CLIMB and other methods. (**E**) Left: Comparison of deconvolution accuracy using Pearson correlation coefficient (PCC) between predicted and true cell line proportions. Statistical significance assessed using Pearson and Filon method with *cocor* R package. Left boxplot depicts PCC *per sample* (4 true versus 4 predicted cell line abundances for 6 bulk samples). Right: same approach using Spearman Correlation Coefficient. (**F**) Left: Similar to (**E**) but for R^2^ metric. Right: similar to (**E**) but for root mean square error (RMSE) metric.

### Evaluating Potential Bias Towards Cellular Proportions Contained In Reference Dataset

The deconvolution process of CLIMB, which employs the entire single-cell matrix as input, may introduce bias mirroring the cellular proportions in the reference matrix. To mitigate such bias, CLIMB’s model executes three sequential deconvolutions. After deconvoluting cellular proportions during its first iteration, CLIMB subsets cells in the reference matrix based on these proportions, which subsequently serve as a reference for the next deconvolution. This process is iterated twice, progressively adjusting the reference matrix to align its cellular proportions with those of the target bulk sample (SFig. 2, Methods). To demonstrate CLIMB’s model accuracy and robustness, we studied how the correlation between cellular proportions in the reference matrix and target bulk samples impacts deconvolution accuracy. We simulated eleven scenarios ranging from strong to zero-correlation contexts, using reference and pseudo-bulk data derived from AML cells in the Van Galen et al. dataset (SFig. 3A). We examined accuracy and resilience to induced bias, which we inferred from the variability in accuracy across our range of bias contexts. By comparing CLIMB (coined ‘CLIMB-corrected’) with the first iteration of CLIMB algorithm (‘CLIMB-init’) to a panel of seven deconvolution methods, we observed that CLIMB was the second most accurate method, and the third less biased method (SFig. 3B-D). Interestingly, CLIMB demonstrated enhanced accuracy and resilience to bias over CLIMB-init in most cross-dataset pseudo-bulk analysis, particularly in cases with significant cellular proportion imbalances between reference and target bulk (SFig. 3E-F). Overall, we demonstrate that CLIMB iterative deconvolution exhibits high accuracy while maintaining low bias.

### Benchmarking Bulk Deconvolution Methods using scRNA-seq Derived Pseudo-bulk Samples in a Cross-Dataset Context

To extend our in vitro findings, we conducted cross-dataset deconvolution by integrating five pairs of scRNA-seq datasets in five cancer types, totaling ten cross-dataset analyses [24, 41–48]. These cross-dataset analyses first involved transferring cell-subtype labels between datasets. Then, one dataset served as the reference single-cell dataset while the other was used to generate pseudo-bulk samples (Fig. 3A). Importantly, the reference dataset and the pseudo-bulk datasets were constructed using only non-batch-corrected raw counts to preserve a realistic batch effect across datasets. The comprehensive results of this large-scale analysis demonstrated that CLIMB significantly outperformed other deconvolution methods (Fig. 3B), based on an aggregated ranking across four evaluation metrics (PCC, SCC, RMSE, and R2). In the context of Acute Myeloid Leukemia (AML), CLIMB outperformed other methods (Fig. 3C) in (i) ‘Van Galen > Naldini’ deconvolution, with CLIMB achieving the best PCC, SCC, and R2 (Fig. 3D, F, SFig. 4E), (ii) ‘Naldini > Van Galen’ deconvolution, where CLIMB achieved the highest PCC, RMSE, and R2 (Fig. 3E, F, SFig. 4G), and (iii) ‘200 simulated pseudo-bulks’, a dataset of 200 simulated bulk samples generated from the Van Galen et al. dataset, where CLIMB demonstrated the top scores for all four metrics (SFig. 4F). Interestingly, CLIMB demonstrated superior performance in predicting the proportion of leukemic stem cells (‘HSC-like’ in Van Galen dataset) in both analysis with available HSC-like labels (Fig. 3G). Moreover, CLIMB ranked first for CRC cancer analysis (SFig. 4H-J). CiberSortX and CLIMB emerged as the best methods in Melanoma analysis (‘Tirosh > Jerby’ and ‘Jerby > Tirosh’, SFig. 4K-M). In the Glioblastoma context, CLIMB was the top-performing method both in the ‘10x > SmartSeq2’ and the ‘SmartSeq2 > 10x’ analysis (SFig. 4N-P). In breast cancer context, MuSiC performed best, based on ‘Gray > Wu’ and ‘Wu > Gray’ analyses (SFig. 4Q-S). Overall, these findings highlight CLIMB’s top-ranking accuracy in cross-dataset pseudo-bulk analyses, especially in AML and Glioblastoma contexts, with strong evidence of high accuracy in estimating the HSC-like cell subtype population.

**Fig. 3.**
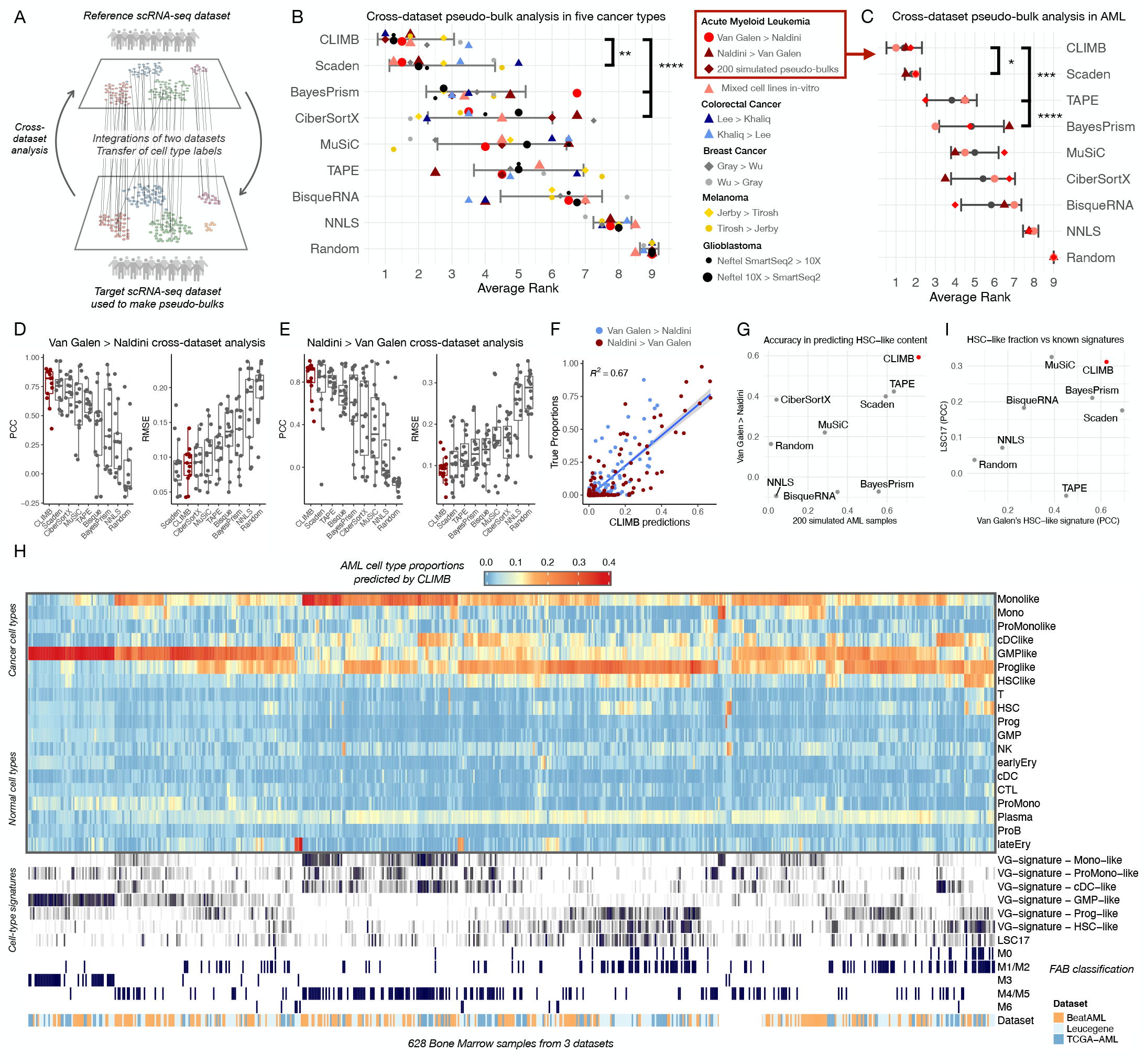
Cell Subtype Composition Deconvolution in Acute Myeloid Leukemia. (**A**) Experimental design outline for cross-dataset deconvolution analysis. (**B**) Average rank across four metrics (PCC, SCC, R2, and RMSE) for 10 cross-dataset pseudo-bulk deconvolutions, including in-vitro mixes and 200 simulated AML samples, totaling 12 analyses. (**C**) Subset of (**B**), focusing on the four AML context analyses. (**D**) Pseudo-bulk analysis using Van Galen dataset as a reference for Naldini’s pseudo-bulks. Boxplots depict PCC/RMSE values for each pseudo-bulk sample (13 samples, 19 cell subtypes’ predictions). (**E**) Similar to (**D**), but in the Naldini > Van Galen context, with 16 data points for the 16 samples in the Van Galen dataset. (**F**) Scatter plot comparing true proportions with deconvoluted proportions using CLIMB (left) and Scaden (right) for both Van Galen > Naldini and Naldini > Van Galen analyses. (**G**) Predictive accuracy of HSC-like cell subtype proportions for Van Galen > Naldini analysis, comparing 13 predicted proportions with true proportions. (**H**) Inferred cell-subtype proportions using CLIMB for bone marrow in BeatAML, Leucegene, and TCGA-AML cohorts (N=628), along with scores of seven signatures (LSC17 and six Van Galen signatures) and FAB labels.

### Bulk Deconvolution of Cell Subtype Abundance Across Three AML Patient Cohorts

We performed deconvolution of cell subtype proportions using bulk RNA-seq samples from three AML cohorts: beatAML [7], Leucegene [49], and TCGA-AML [50], using as a reference only cells in pre-treatment conditions from Van Galen dataset [24]. Employing CLIMB, we delineated a landscape of cell subtype proportions in AML (Fig. 3H). To assess deconvolution accuracy in patient samples, we juxtaposed CLIMB-derived proportions with FAB classification [51], a scheme categorizing patients according to blast (cancer cell) differentiation patterns. Specifically, blasts from M0/M1 patients, characterized by minimal differentiation, were expected to correlate with higher proportions of HSC-like and/or Progenitor-like cancer cells, a correspondence verified by CLIMB-derived proportions (Fig. 3H, SFig. 5A). Similarly, M3 samples (PML-RARA classified patients) were anticipated to exhibit enrichment for GMP-like cells, in line with our deconvoluted GMP-like proportions (SFig. 5A). Moreover, we observed monocyte-like cancer cells to be enriched in M4/M5 patients, along with erythrocyte enrichment in M7 patients, both of which were expected (SFig. 5A). To evaluate CLIMB’s accuracy in predicting leukemic stem cells, we correlated deconvoluted HSC-like proportions with the LSC17 signature [31]. As expected, the highest correlations were observed with Prog-like and HSC-like cell subtypes (SFig. 5B). Interestingly, proportions of HSC-like cells predicted by CLIMB showed a noticeable alignment with both the HSC-like signature established by Van Galen et al. [24] (SFig. 5C) and the LSC17 signature, when compared to other deconvolution methods (Fig. 3I). To evaluate the consistency of results across different deconvolution algorithms, we assessed cross-method correlations across all three AML cohorts. We observed that the top three methods—CLIMB, Scaden, and CiberSortX—formed a distinct cluster (SFig. 5D). In summation, CLIMB provides a robust depiction of the cellular landscape of AML patients across three cohorts, laying foundations for deconvoluting cell subtype drug sensitivity.

### Benchmarking Bulk Expression Deconvolution into Cell Subtype Expression and Simulated Over-Expression

CLIMB enables the deconvolution of bulk-level RNA expression into cell subtype-specific expression. This can be achieved through either an ‘overall’ approach, which accounts for expression averaged across all bulks used as input, or a ‘high-resolution’ mode that disentangles cell subtype expression within each bulk sample. In our in-vitro experiment with leukemic cell lines, CLIMB’s ‘overall’ deconvoluted expression achieved high accuracy when compared to bulk RNA-seq expression obtained from pure cell lines (PCC of 0.94, SFig. 6A-C). BayesPrism displayed similar patterns and accuracy to CLIMB, achieving a PCC of 0.92 (SFig. 6C). Subsequently, we deconvoluted AML cell-subtype expression based on the ‘Van Galen > Naldini’ cross-dataset framework. Notably, the ‘overall’ mode of CLIMB demonstrated compelling deconvoluted expression patterns for cancer cell markers (PCC of 0.75, SFig. 6D-F), surpassing the performance of BayesPrism (0.59) and CiberSortX in ‘group mode’. We then subjected CLIMB’s deconvoluted expression to a ‘high-resolution’ test, simulating the over-expression of 20 genes in the HSC-like subtype. This induction was applied to 7 out of 13 pseudo-bulk samples, with varying fold changes. The simulated expression was then compared to true expression values, showing CLIMB’s deconvoluted expression high accuracy at fold change 10 and higher (SFig. 6G). To assess the precision in detecting differentially expressed genes (DEGs), we conducted a DE analysis between wild-type (WT) and over-expression (OE) conditions. We employed the AUC-ROC metric to evaluate DEG detection accuracy (SFig. 6J-K). BayesPrism exhibited the highest accuracy in recovering DEGs followed by CLIMB (SFig. 6J). As a positive control, we checked that true simulated expression efficiently recovered DEGs (AUC > 0.95 with fold change > 2, SFig. 6J, K). We then computed correlations (PCC) between true and deconvoluted expression (SFig. 6L), showing BayesPrism as the most accurate (median PCC of 0.76), followed by CLIMB (0.66). In summary, CLIMB demonstrates accuracy in expression deconvolution, particularly in ‘overall’ mode.

### Benchmarking Drug Sensitivity Deconvolution with In-Vitro Cell Line Mixtures

In this next step, we integrate drug sensitivity data (measured as a cell viability rate) with CLIMB’s bulk deconvolution results. To achieve this, we developed CLIFF, a method that deconvolutes bulk-level cell viability into cell-subtype-specific cell viability (Fig. 1, 4A). We generated drug sensitivity data from *in vitro* mixtures comprising six combinations of four human leukemia cell lines, as previously depicted in Fig. 2 (HL60, SUDHL4, K562, and THP1). These mixtures were exposed to seven drugs commonly employed in AML consolidation therapies (5-Aza-2’-deoxycytidine, cytarabine, imatinib, nilotinib, doxorubicin, rapamycin, mitomycin C). To assess the accuracy of drug sensitivity deconvolution, we determined the ground truth cell viability of the four individual cell lines when exposed individually to the same drug panel, doses, and time points. We initially evaluated the theoretical basis of CLIFF’s model by reconstructing the experimentally-assessed cell mix viability from the experimentally-determined viability of pure cell lines. This was achieved using CLIFF’s model equations (Equations 15, and 16) to correlate both metrics (SFig. 7A), which showed a high Pearson’s correlation of 0.88. In the next step, we evaluated CLIFF’s deconvoluted cell viabilities against ground truth, showing a PCC of 0.80 across all 21 conditions (Fig. 4B, C). When utilizing high-resolution cell-subtype expression as input (‘CLIFF-highres’), the accuracy of CLIFF was diminished (PCC of 0.59, Fig. 4C, SFig. 7B). Subsequently, we conducted a comparison of CLIFF’s accuracy with the approach employed by Zeng et al. [34], which relies on a correlation between cell subtype proportions and bulk-level drug sensitivity; with Bottomly et al.’s approach [32], wherein the PC1 of cell subtype signatures is correlated with bulk drug sensitivity; and a linear regression using CLIMB proportions as input and bulk-level cell viability as ouput (LinReg). When assessing drugs and timepoints individually, CLIFF exhibited the highest correlation (median PCC of 0.82, Fig. 4D), followed by Bottomly (0.80), ‘CLIFF-highres’ (0.68), Zeng (0.68), and LinReg (0.51). Interestingly, CLIFF is the only method that predicts cell viability rather than providing a drug sensitivity score based on correlation metrics, which results in a significantly lower RMSE compared to all other methods, both in ‘overall’ and ‘high-resolution’ modes (Fig. 4E). Thus, within a framework comprising six cell mixes encompassing four cell lines, CLIFF exhibited precise deconvolution of drug sensitivity and proved effective to integrate cell subtype proportions estimated via CLIMB.

**Fig. 4.**
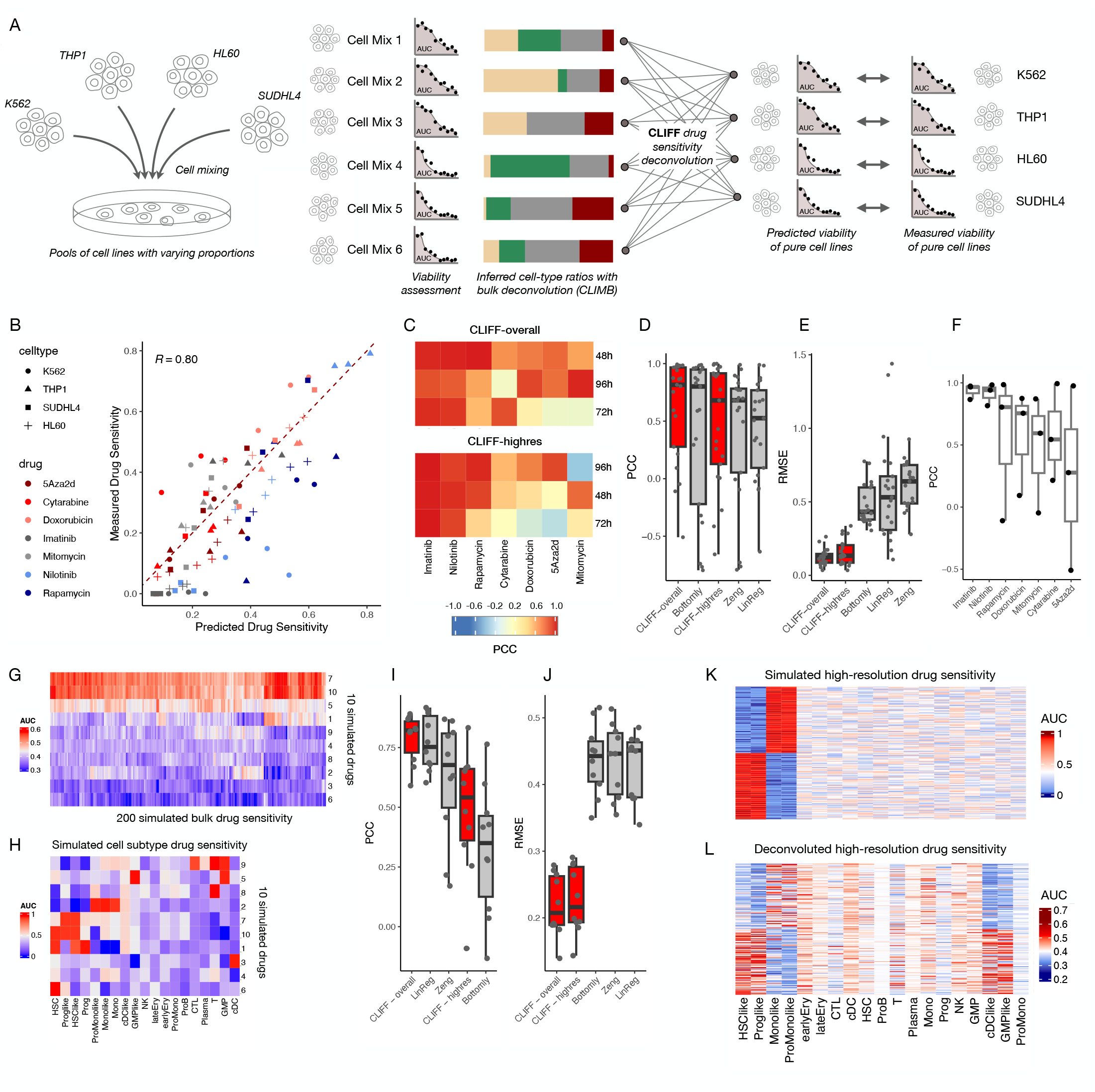
Deconvolving Cell Line Drug Sensitivity: *In-Vitro* and Simulated AML Context. (**A**) Experimental setup: Known cell line mixtures are deconvoluted to estimate cell abundances, which are integrated with bulk drug sensitivity for predicting individual cell line drug responses. (**B**) Predicted vs. actual cell line drug sensitivity for multiple timepoints, cell lines, and drugs. (**C**) Heatmap displays predictive accuracy as PCCs for each drug and timepoint. (**D**) Method comparison of deconvolution accuracy using PCCs. (**E**) Method comparison of deconvolution accuracy using RMSE values. (**F**) CLIFF’s predictive accuracy as PCC, grouped by drugs. (**G**) Drug sensitivity simulation on 200 pseudo-bulk samples derived from cell subtype sensitivities (**H**). (**H**) Simulated cell subtype drug sensitivity pattern for 10 drugs, used for generating bulk drug sensitivity. (**I**) Predictive accuracy as PCC for CLIFF and other methods, deconvoluting cell subtype sensitivity (**G**) using bulk drug sensitivity (**H**). (**J**) Same as (**I**), but using RMSE metric. (**K**) Simulated high-resolution drug sensitivity across pseudo-bulk samples and cell subtypes. (**L**) Deconvoluted high-resolution drug sensitivity based on bulk samples from (**K**).

### Simulated *Ex-Vivo* Drug Sensitivity Assay for Benchmarking Deconvolution of Drug Sensitivity

We then simulated bone marrow samples from AML patients to generate a benchmarking dataset similar to the beatAML cohort; as the latter lacks ground truth data for testing deconvolution. Using the Van Galen single-cell dataset, we generated 200 pseudo-bulk RNA-seq samples along with their corresponding bulk cell viability data for ten simulated drugs (Fig. 4G). To do this, we simulated cell viability for 19 cell subtypes across ten drugs (Fig. 4H), which serves as our ground truth for evaluating CLIFF’s deconvolution accuracy. We proceeded to assess the accuracy of drug sensitivity deconvolution using the same five approaches mentioned earlier (Fig. 4D-E). In this context, CLIFF-overall demonstrated the highest accuracy, with a median PCC of 0.82 (Fig. 4I), followed by LinReg (0.75), Zeng (0.68), CLIFF-highres (0.54), and Bottomly (0.35). As observed in the in-vitro experiment, CLIFF’s advantage, both in overall and high-resolution modes, lies in generating realistic cell viability values, which translates into significantly lower RMSE values compared to every other methods (Fig. 4J). Investigating the effect of the number of input genes on CLIFF’s performance, we found that CLIFF-overall maintained consistent accuracy regardless of the gene subset (SFig. 7E), while CLIFF-highres excelled particularly with higher gene counts (>500). Additionally, introducing L2 regularization improved CLIFF-highres predictions, although it had no significant impact on CLIFF-overall predictions (SFig. 7E). The impact of input size on CLIFF’s accuracy was examined by varying the number of bulk samples (SFig. 7E). Notably, approximately 25 cell mixes proved sufficient for CLIFF to achieve a PCC > 0.6 when deconvoluting 19Acell subtypes, and about 75 input mixtures were required to attain a PCC > 0.7. By simulating patient-specific mutations (SFig. 7F) that were associated with heightened drug sensitivity in HSC-like/Prog-like cell subtypes and lowered sensitivity in Mono-like/ProMono-like cell subtypes, we examined how well CLIFF-highres could discern patient-level variability (Fig. 4K, L). Our observations showed that CLIFF-highres detected shifts in drug sensitivity between the two sample groups in the correct cell subtypes (HSC-like, Prog-like, Mono-like, and ProMono-like). However, it also introduced false positive predictions in cDC-like and GMP-like subtypes. In summary, based on our benchmarking of CLIFF in both in-vitro experiments and simulated contexts, it appears that CLIFF’s ‘overall’ mode effectively predicts averaged cell-subtype drug sensitivity. Therefore, we will limit the use of CLIFF to its ‘overall’ mode in the next steps.

### Deconvoluting Bulk-Level Drug Sensitivity to Cell Subtype Sensitivity Using beatAML Cohort

Utilizing cell subtype proportions and expression data estimated by CLIMB from the beatAML cohort, we integrated this information with bulk *ex-vivo* drug sensitivity data using CLIFF, which revealed a heterogeneous landscape of drug sensitivity patterns among distinct cell subtypes (Fig. 5A). We assessed the findings based on 14 studies that documented cell-subtype level drug sensitivity and featured 22 pairs of drug-to-cell subtype interactions (Supplementary Tab. 1). CLIFF’s predictions aligned with existing literature for venetoclax [12, 13], erlotinib [16], A-674563 [17], AZD1480 [18, 19], and crenolanib [20], indicating elevated sensitivity in HSC-like and Prog-like cancer cells. Interestingly, midostaurin [21] and selinexor [22] were predicted to target HSC-like cell subtypes, but not progenitor cancer cells (Prog-like). We also documented drug sources that targeted differentiated blasts. Doramapimod, predicted by CLIFF to target monocyte-like cells, aligns with previous evidence of heightened activity in M4/M5 patients showing expression of CD14 monocyte marker [11]. Similar agreement was observed for trametinib [12] and selumetinib [9], known to target differentiated blasts, and projected by CLIFF to exhibit strong affinity for monocyte-like cancer cells. Bromodomain and Extraterminal Domain Inhibitors (BETi) were also shown by Romine et al. [10] to target differentiated blasts, which coroborates CLIFF’s predictions on BET-inhibitor JQ1, predicted to preferentially target monocyte-like cancer cells. Ruxolitinib [12], however, did not display a clear pattern, as it was predicted to target pro-monocyte-like but not monocyte-like cancer cells. CLIFF’s predictions also indicate a notable resistance of monocyte-like cells to venetoclax, validated by four distinct studies [11–15]. Among the 22 documented instances of cell subtype-specific drug sensitivity, CLIFF demonstrated 16/22 cases with clear concordance (72.7%), 3/22 with partial alignment, and 3/22 in direct contradiction (13.6%). Furthermore, we evaluated the concordance between CLIFF’s predictions and results reported by Bottomly et al. using the same dataset [32], revealing an overall correlation of 0.44 for all drugs (SFig. 8A), with stronger correlations observed for established drug-to-cell subtype associations (SFig. 8B).

**Fig. 5.**
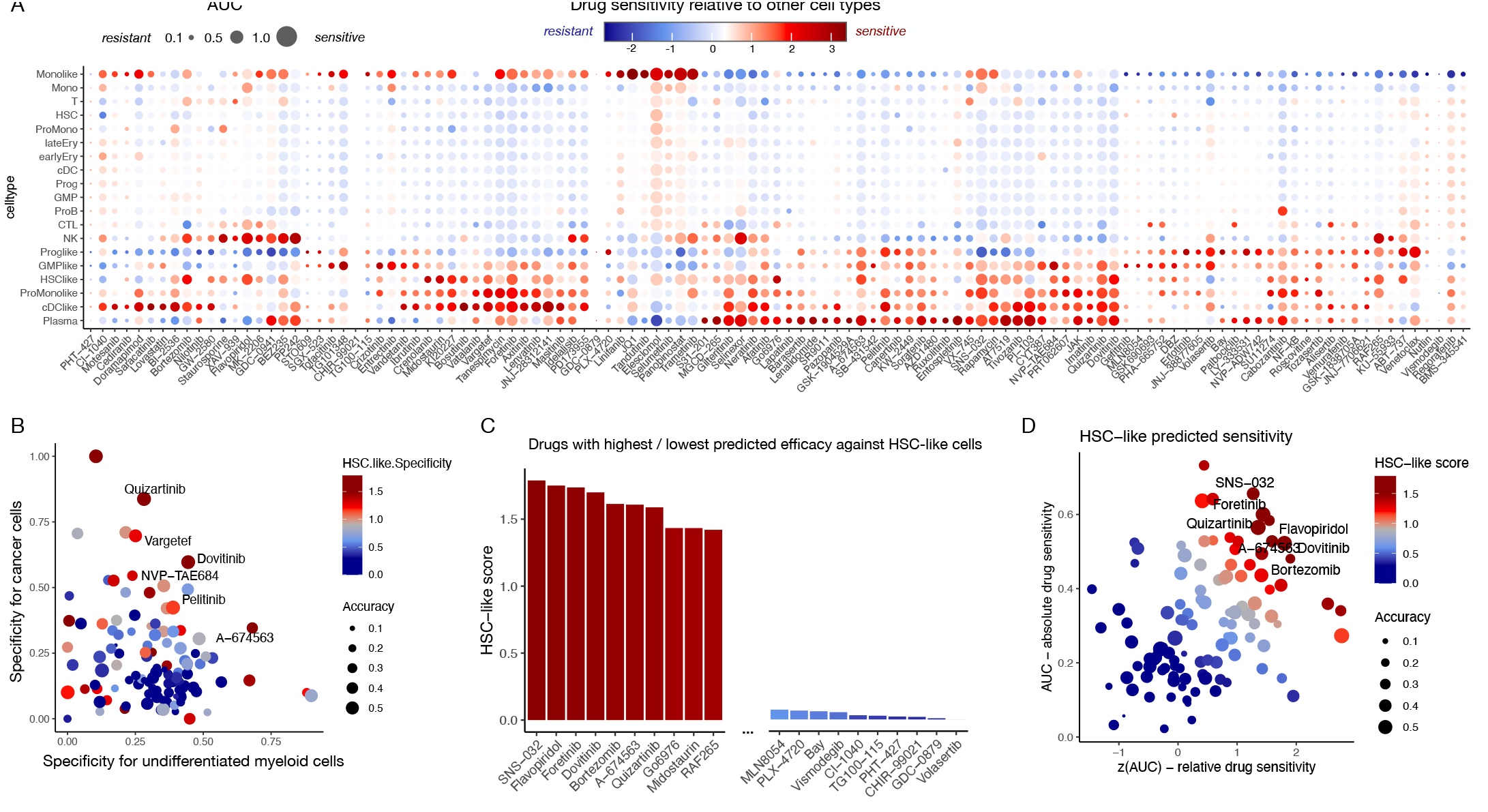
Deconvolving Cell Subtype-Specific Drug Sensitivity in AML *ex-vivo* Data. (**A**) Deconvoluted drug sensitivity across cell subtypes in the beatAML cohort dataset, with dot size indicating AUC of sensitivity and color representing standardized AUC, indicating relative drug sensitivity among cell subtypes for a specific drug. (**B**) Differentiated (Mono/ProMono/cDC) vs. undifferentiated (HSC/Prog/GMP) myeloid cells’ sensitivity (grouped) on the x-axis and normal vs. cancer myeloid cells’ sensitivity on the y-axis. (**C**) HSC-like sensitivity score computed from AUC and standardized AUC. (**D**) Drug sensitivity of HSC-like cells shown on a scatter plot with AUC on the x-axis, standardized AUC on the y-axis, and color indicating the combined score. (**E**) High-resolution drug sensitivity landscape for venetoclax across patients, with boxplots showing distribution among patients, and bottom heatmap displaying somatic mutation presence. (**F**) Similar view as (**E**), but for A-674563 compound.

Subsequently, we focused on evaluating drug sensitivity in HSC-like cells, ranking drugs according to their potential effectiveness against this particular cell subtype (Fig. 5B-D). Among the top ten drugs, only A-674563 and midostaurin had available validation from the literature (ref Tab). The eight undocumented drugs predicted to be the most effective against HSC-like cells were, in order, SNS-032, flavopiridol, foretinib, dovitinib, bortezomib, quizartinib, Go6976, and RAF265. In summary, the combined use of CLIMB and CLIFF successfully unveiled a comprehensive landscape of drug sensitivity at the cell subtype level in AML, along with the ability to predict drug sensitivity in the clinically relevant HSC-like cell subtype.

### Single-Cell Resolution Prediction Identifies Resistant Cancer Cell Subpopulations

We further developed CLIFF-SC, which computes drug sensitivity prediction at single-cell resolution (Fig. 6A, Methods, Supplemental Methods), leveraging coefficients derived by CLIMB for each individual cell. We focused on three drugs predicted to target the HSC-like compartment: venetoclax, A-674563, and foretinib, and one drug with an opposite pattern, nilotinib, which targets monocyte-like cancer cells (Fig. 5A). We visualized single-cell survival probability on 2D projections (UMAP) using all cancer cells from Van Galen dataset [24], which showed considerable heterogeneity in distributions across the four drugs and the six cancer cell subtypes (Fig. 6B-E). We observed high concordance between average cell subtype sensitivity predicted by CLIFF-SC and the one predicted by CLIFF-overall (Fig. 6F). Interestingly, we observed a long upper tail of seemingly resistant HSC-like cells for venetoclax compound (Fig. 6B, right panel, circled in red). We thus decided to study in more detail this population of resistant HSC-like. We performed differential expression analysis between resistant and sensitive HSC-like populations, which were clustered using single-cell venetoclax sensitivity predictions (25% most sensitive versus 25% most resistant, Fig. 6G). We could show a downregulation of NPM1 expression and upregulation of ANXA1 and TMSB4XP8 in the 25% with highest predicted resistance to venetoclax (Fig. 6H). These data suggest that expression of these genes could be predictive of venetoclax sensitivity at single-cell level.

**Fig. 6.**
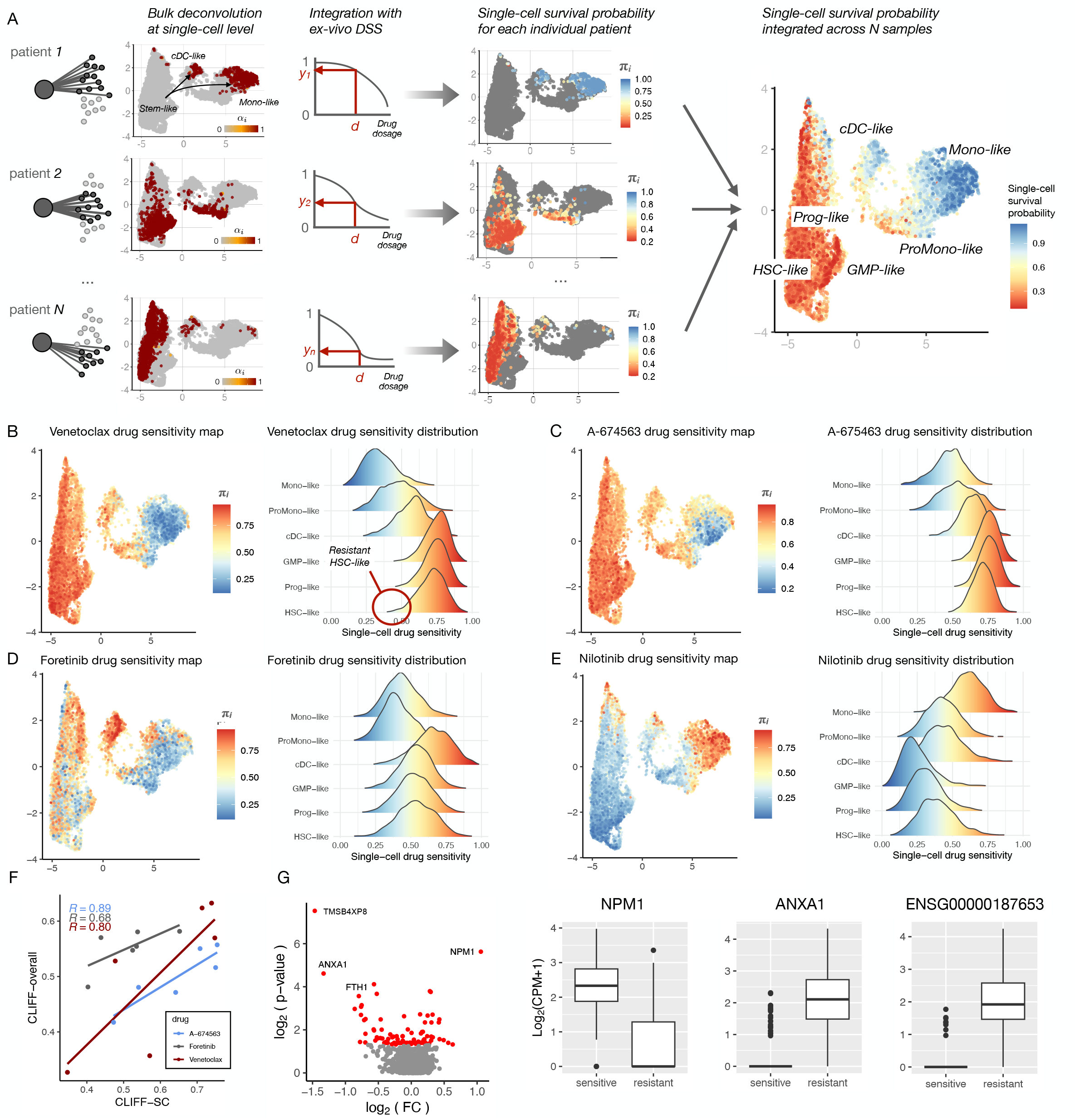
Single-Cell Resolution Drug Sensitivity Prediction. (**A**) Framework overview: CLIMB deconvolution generates coefficients for each single cell (left), integrated with *ex-vivo* drug screening data (DSS) for single-cell survival probabilities (right). (**B**) Single-cell survival probability for venetoclax, based on UMAP projections of cancer cells from Van Galen et al. [24] (left), and survival probability distributions among six cancer cell subtypes (right). (**C**) Similar view for A-674563 single-cell drug sensitivity. (**D**) Similar view for foretinib single-cell drug sensitivity, highlighting upper tail of resistant HSC-like cells in red circle. (**E**) Similar view for nilotinib single-cell drug sensitivity. (**F**) Volcano plot showing gene expression comparison between venetoclax-resistant (25%) and sensitive cells (25%) with p-value versus log fold change. (**H**) Genes over-expressed in venetoclax-resistant population of HSC-like cells.

## DISCUSSION

The beatAML consortium achieved the most extensive *ex-vivo* drug sensitivity screening (DSS) to date across all cancer types [7], paving the way for deciphering cell subtype-specific drug sensitivity. Our study proposes a method to address this challenge, potentially replacing resource-intensive investigations commonly required for assessing cell population-level drug sensitivity in large patient cohorts. We present a pioneering approach to deconvolute bulk drug sensitivity data into its constituent cell subtypes, employing a method capable of retrieving realistic AUC metrics for cell subtype-level drug sensitivity. Through extensive evaluation in cell line mixes and acute myeloid leukemia (AML), we predict drug sensitivity for 19 cell subtypes, 115 anti-cancer drugs, and 286 patients, culminating in a comprehensive dataset comprising over 344,318 data points derived from the beatAML dataset.

Bulk deconvolution is a cost-effective method to uncover cell composition in patient cohorts. To this end, we introduce CLIMB, a novel algorithm that outperforms current standards in various contexts, achieving accuracy comparable to deep learning-based methods with a seemingly simple linear model. CLIMB’s advantage lies in its use of the complete single-cell expression matrix, unlike other methods that uses average cell subtype expression across bulk samples. This is crucial, as cancer cell expression variability among samples is expected. Based on or cross-dataset pseudo-bulk study, CLIMB seems to effectively address batch effects. Notably, CLIMB excels in estimating proportions of HSC-like cells, a subtype linked to poor AML prognosis and relapse, enhancing predictions of their drug sensitivity. We anticipate CLIMB’s utility across diverse biological scenarios, particularly in deciphering cancer sample cellularity.

By considering latent variables and employing an Expectation-Maximization algorithm, CLIFF deconvolves bulk-level cell viability into its sub-population-level components (Method, Supplemental Methods). We propose that the primary strength of the CLIFF method, when compared to prior methodologies [32–34], lies in its capacity to generate biologically interpretable metrics of cell viability at the cellular level. This stands in contrast to correlative techniques that merely provide a score linking cell subtypes and drugs without delivering a meaningful biological measure. We demonstrated CLIFF abilities first with engineered mixes of leukemic cell lines and pseudo-bulk analysis. We explored multiple avenues for extending CLIFF’s ‘overall’ mode—which provides an averaged prediction across all bulk samples taken as input—with two higher resolution strategies. On one hand, CLIFF-highres, which uses cell subtype expression and mutation data as input, delivering cell viability predictions for each sample. In our benchmarking, CLIFF-highres generally showed decreased accuracy compared to CLIFF-overall, likely due to the limited number of input samples. In contrast, CLIFF-SC employs a full-sized single-cell dataset as input, substituting cell subtypes with single cells in CLIFF’s original model, thereby allowing the prediction of single-cell drug sensitivity. It proved to be a straightforward method to enhance CLIFF’s resolution, providing valuable single-cell maps of drug sensitivity. We then turned to the beatAML dataset, using CLIFF-overall model enhanced with mutation data as covariates. In this setting, we could reveal an extensive landscape of cell subtype drug sensitivity. Also, we predicted A-674563, Midostaurin, SNS-032, and foretinib, among other compounds, to be efficient at targeting HSC-like cells. We validated our findings by relying on 14 publications that documented cell-subtype level drug sensitivity through functionnal experiments in AML patient’s samples, for 22 pairs of drug to cell subtype interactions [9, 11–13, 16–23], with showed overall good agreement with our predictions (72.1%).

Our approach has some limitations. First, since ground truth cell viability data is not available for patient samples, it is challenging to assess our deconvolution results on AML cohorts. This limitation led us to use a small compilation of literature as a comparison point. Second, any weakness in CLIMB’s model could potentially affect CLIFF’s predictive accuracy. We believe CLIFF would benefit significantly from being used in contexts where cellular proportions can be directly observed and measured rather than predicted through deconvolution. For example, using a cohort of patients with available scRNA-seq data and cell viability measures. Third, we rely on cell viability observed ex-vivo from bone marrow samples in 96-well cell viability plates, an environment which might bias the viability of fragile cell subtypes.

By integrating diverse data modalities, our combined approach seeks to enhance cellular composition predictions and unravel drug sensitivity at the level of individual cell subpopulations. Here, we make a pioneering effort, as far as our knowledge extends, to advance drug sensitivity deconvolution to novel resolutions, encompassing personalized and single-cell-level predictions. As the integration of single-cell RNA-seq and ex-vivo drug sensitivity assays gains traction for diagnostic applications, the development of suitable computational methodologies becomes imperative to fully leverage the wealth of information from these joint data sources.

## METHODS

### Gene and cell selection

In our approach to gene subset selection prior to bulk deconvolution, we utilized the Scran::ModelGenVar function on the expression matrix derived from single-cell RNA sequencing (scRNA-seq) data. By applying this methodology, we identified genes with estimated biological variance surpassing technical variance, as indicated by positive biological variance outcomes from ModelGenVar. This strategy eliminated the reliance on arbitrary thresholds for the determination of variable or deferentially expressed genes and was consistently employed across all analyses in this study.

Regarding cell selection, in the context of our in-vitro experiment, we observed cells forming five clusters, with four clusters being distinctly attributed to the four mixed cell lines. The fifth cluster lacked a clear marker expression pattern, exhibited smaller size (only few cells) compared to the other clusters, and displayed elevated mitochondrial read content. Consequently, we excluded this ambiguous cluster from our analysis, while retaining the remaining cells in four clusters that were attributed their known cell line labels according to their marker expression pattern.

Regarding the re-analysis of publicly available scRNA-seq datasets, we harnessed the cell selection information provided in the processed data tables, abstaining from further cell filtering based on specific criteria. In instances where datasets encompassed over 30,000 cells, we subsampled them to reach a count of 30,000 cells, adopting the set.seed(1) option for reproducibility. This enabled us to execute the Seurat integration tool without memory constraints. The datasets subjected to subsampling included Naldini et al., Gray et al., Wu et al., and Khaliq et al.

### Prototypical bulk deconvolution model

The goal of a bulk deconvolution model is to fit coefficients w_k_ corresponding to each cell-subtype k through a linear model. Fitted coefficients directly relates to cell-subtype proportions. Thus, bulk deconvolution solves the following problem :

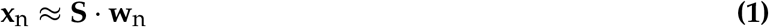

with **x**_n_ representing a vector of gene expression of length G, for a given mixture n. The matrix **S** is a G by K matrix containing the average expression for each cell-subtype k and each gene g. To obtain proportions w_nk_ that sum up to 1, the model can either impose a constraint Σ_k_ w_nk_ = 1, or the coefficients w_nk_ can be normalized after fitting. In this model, the genes are our observations, and corresponds to genes shared between **x**_n_ and **S** or to a subset of marker genes. Note that every mixture n are resolved with an independent linear model, and that the average expression of cell-subtypes is usually derived from a scRNA-seq that does not contain mixture n. Thus, we assume that **S** ≈ **S**_n_. Of note, this simple model do not apply for advanced deconvolution algorithms like CiberSortX, BayesPrism, Scaden, BLADE, or TAPE.

### CLIMB: CELL-LEVEL LINEAR MODEL THROUGH BULK-TO-SINGLE-CELL DECONVOLUTION

We developed the CLIMB method, which proposes an innovative approach to solve a bulk deconvolution problem, by applying a mapping from each bulk sample to each single-cell from a reference dataset. We thus fit the coefficients *α*_i_ for each single-cell i in an intial run of CLIMB, and thus we find the best linear combination of single-cell c_i_ to predict bulk expression **x**_n_. Thus, we use the relation **x**_n_ ∼ **C** *·* ***α***_n_, shown below in more details :

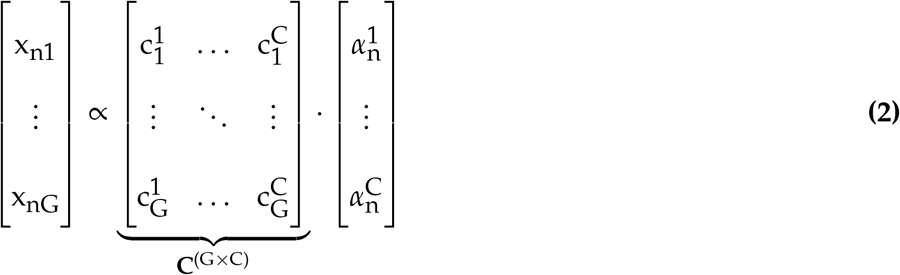

where the vector **x**_n_ represent the bulk expression matrix for a mixture n, **C** is the raw counts matrix from scRNA-seq, and ***α***_n_ is the vector of coefficients fitted for each single-cell i for a given mixture n. Thus, CLIMB loss function is defined by :

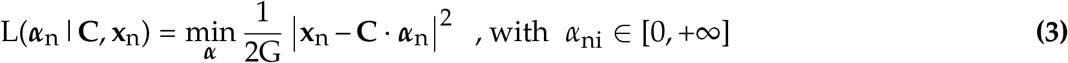

Thus, we impose a non-negativity constraint on the coefficients, and no regularization. We optimize *α*_i_ coefficients associated with each single-cell i with glmnet R library [52], with the options lower.limits = 0, lambda = 0, and scale = TRUE, intersect=FALSE. We indeed observed no improvements of the deconvolution with neither l_1_ nor l_2_ regularization.

We then group and normalize coefficients *α*_ni_ obtained with (3) to obtain the cell-subtype proportions w_nk_ for each cell-subtype k and mixture n :

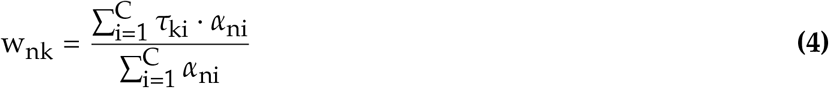

With ***τ***_k_ being an indicator variable of length C, equal to 1 whenever cell i is of type k.

### Empirical Bayes Sub-Sampling Mitigates Bias Induced by Reference Matrix

The equations 2 3, 4 yield biased estimates of cellular proportions towards the cellular proportions contained in the reference single-cell matrix **C**. This bias acts as a prior, impacting CLIMB’s predictions. To enhance cellular composition depiction, we’ll correct **C** by informed cell sub-sampling, creating a new matrix via an Empirical Bayes approach estimating the prior from the data.

We use the initial CLIMB-derived cellular proportions from equations 2 3, 4 to sub-sample half of the cells in reference matrix C based on our predicted biased proportions. Each sub-sample ensures a minimum of 50 cells (or 2% of total cell count) per cell type. In total, five sub-sampling are done, and this process is repeated three times. Ultimately, the proportions from the second iteration - and thus third devoncolution - become the final CLIMB output (stored in ‘props.corrected’). CLIMB’s iterative algorithm is summarized in SFig.2.

Additionally, a gene selection aids CLIMB’s deconvolution. First, dataset-specific top-variable genes are selected based on cell type average expression per gene. Genes with significant fold change compared to the average expression and compared to the second top-expressed cell type, are chosen (threshold set to 95% and 75% fold changes quantiles). Cell types with top expression are recorded, allowing a maximum of 100 genes per cell type. This initial gene selection is followed by bulk sample-specific gene selection: Genes are ranked according to their maximum cell type expression multiplied by their expression in the target bulk sample, and the top 500 genes are selected. These gene selection are thus bulk sample-specific.

### Deconvolution of cell-subtype expression

CLIMB allows the prediction of cancer cell expression in a two step manner. Let first define our target *high-resolution* expression 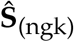, which is the cell-subtype-specific and bulk sample-specific expression for each cell-subtype k. We assume that for a given group of samples, the *grouped* expression 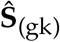 relates to the *high-resolution* expression with the following formula:

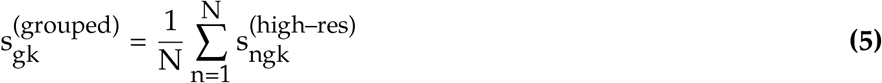

First, we take advantage of the bulk to single-cell mapping to directly derive the cell-subtype expression from the bulk-to-single-cell deconvolution or *mapping*. We can simply sum up the expression of cells mapped by CLIMB to get a *high-resolution* cell-subtype-specific and bulk-specific expression 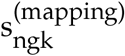.

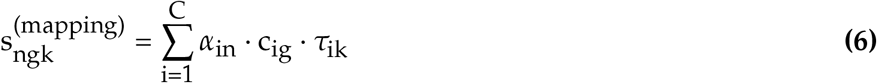

with *α*_in_ being the coefficient fitted by GLMNET on each single-cell, linking each cell i to each bulk n. c_ig_ denotes the single-cell expression of cell i and gene g, and *τ*_ik_ is an indicator vector equal to 1 if cell i is of type k.

Although this is potentially sufficient in cases where scRNA-seq reference dataset maps all cell states present in bulk samples, it can be insufficient to detect genes deferentially-expressed (DE) at cell-subtype level. As DE genes will likely induce error on the cell-subtype expression deconvolution, we use the error itself to fit our model. We can also make the reasonable assumption that DE genes only originates in cancer cells. Thus, for a given gene g and a given bulk n, we have:

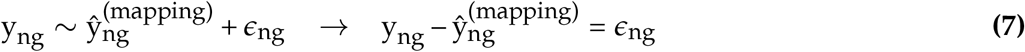

Thus, we use 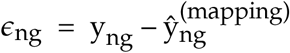 as input for a model to predict cancer cell-subtype expression. We decompose our error *ϵ*_ng_ between the error originated from DE genes 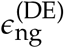, and an additional error term 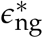 Similar as other methods, we use the information from all bulk samples N to deconvolute the high-resolution cell-subtype specific expression 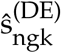 We consider that 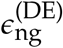 only originates from cancer cells C^(C)^ of type K^(C)^, thus

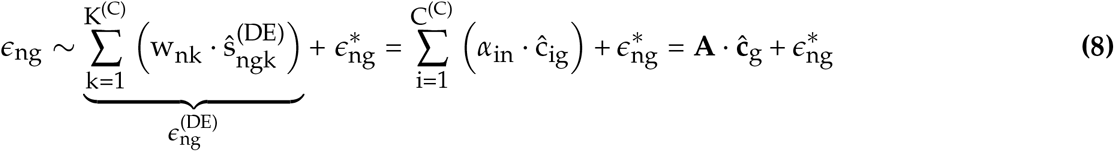

Where ĉ_g_ is the vector of coefficients we wish to predict. We used Glmnet with options lower.limits = 0, lambda = 0, and scale = TRUE, fitting the following loss function:

Our final prediction of cell-subtype expression for a cancer cell-subtype k, a given bulk n, and a given gene g is:

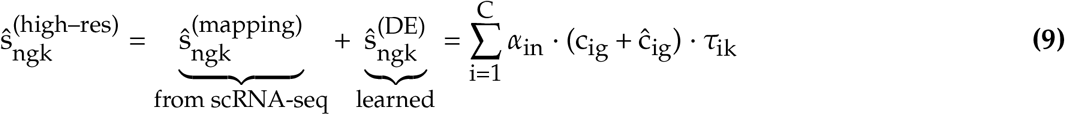

Following the assumptions made above, we set ĉ_ig_ to zero for normal cells, which imply that 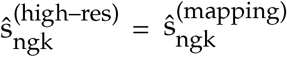 for normal cell subtypes.

### CLIFF - Cell fate inference for drug sensitivity deconvolution

Here we propose to infer the drug sensitivity of individual cell subtypes using only drug sensitivity data observed on mixed cell populations, considering K different cell populations in N mixed samples, with proportions than can be defined as **w**_n1_ = [w_n1_, …, w_nK_], with w_nk_ ∈ [0, 1] and 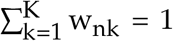. Drug sensitivity observed in the bulk samples we wish to demix are represented by a vector **y**_(N)_, with y_n_ ∈ [0, 1] being the empirical surviving rate of cells, and for sample n. Therefore the target matrix we wish to predict is **Ŷ** ^N*×*D^, and the predicted values of drug sensitivity is assumed to depend on the underlying drug sensitivity of cell subtypes *π*_nk_ ∈ [0, 1] for patient n, and cell-subtype k, represented as a surviving rate of cells.

To model drug sensitivity, we assume that single-cell survival is represented as a binary output *δ*_ni_ ∈ {0, 1} that follows a Bernoulli distribution. Thus, every single-cell i in patient n exposed to a drug can either survive (*δ*_ni_ = 1) or die (*δ*_ni_ = 0). Thus a cell i has a probability P(*δ*_ni_ = 1) to survive and 1 – P(*δ*_ni_ = 1) to die. Let’s say we are interested to know the survival probability P(*δ*_ni_ = 1), given a drug dose. We could resolve the log-likelihood of the Bernoulli distribution for all the cells of a single sample n :

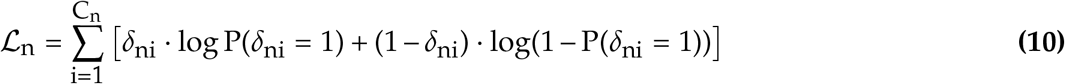

It turns out that in our problem, *δ*_ni_ cannot be directly observed. We are indeed only able to observe the overall ratio of cells that survives among all cells C_n_ for a given patient n, which we define as y_n_. It relates to the values *δ*_ni_ with :

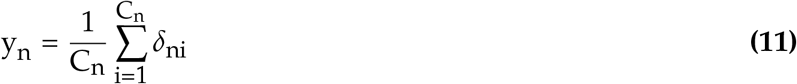

Thus, by multiplying (10) by 1/C_n_, if we assume that every cell i share the same average probability P(*δ*_nid_ = 1), we can re-write (10) and obtain the following log-likelihood :

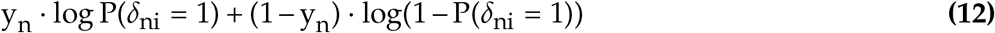

As stated above, the core of the problem is to assume different survival probability for our single-cells. To slightly simplify this, we can consider a cell-subtype-specific survival probability. Thus, we consider the survival probability observed in our mixed population of cells P(*δ*_nid_ = 1), as a mixture of K probabilities 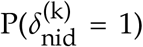, one for each cell subtype. Let introduce *π*_kn_, a variable describing the survival probability of cell-subtype k:

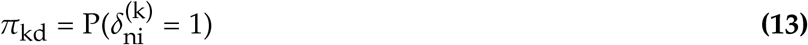

We also define *v*_nd_, which corresponds to the expected number of cells surviving in patient n with drug dose d, and assume it follows a Binomial distribution, with y_nd_ being the ratio of surviving cells among cells in C_n_. We further define Ŷ_nd_ as an average of cell-subtype survival probabilities *π*_kdn_ constituting the bulk, weighted with cell-subtype proportions w_nk_:

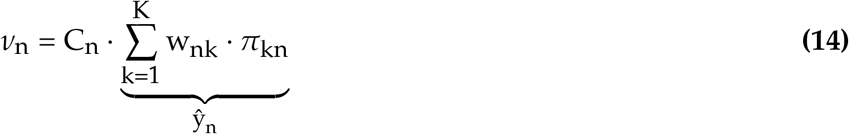

with w_nk_ the cell-subtype proportions. Ŷ_n_ is the predicted cell surviving rate in the sample n, and thus the target we wish to predict through a cross-validation procedure. We can say that if we let :

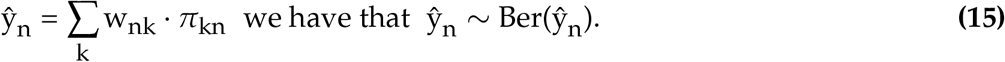

Additionally, we impose the cell-subtype-specific survival probabilities *π*_kdn_ to be non-linearly related with cell-subtype gene expression **s**_kn_ (a vector of length G) through a sigmoid function:

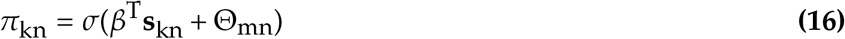

Note that the sigmoid in (16) also optionaly depends on Θ_mn_, which is an intersect associated with each mutation m in each patient n, attributed to the value 1 in each cancer cell type and to 0 in normal cell types. To clarify the notation, we set that 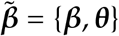, with ***θ*** being a vector containing coefficients associated with each mutation m.

When running CLIFF in *overall* mode, there are two assumptions that we use to simplify the problem. First, we assume that every sample n share the same cell-subtype expression s_gk_, for a given gene g and cell-subtype k. Thus, s_gk_ ≈ E_N_[s_gkn_]. Second, we assume that cells of the same cell-subtypes across the N samples share the same sensitivities *π*_k_. We thus assume that *π*_k_ correspond to the expected drug sensitivity over N samples, and therefore that *π*_k_ ≈ E_N_[*π*_kn_].

Thus, the overall log-likelihood for our problem is :

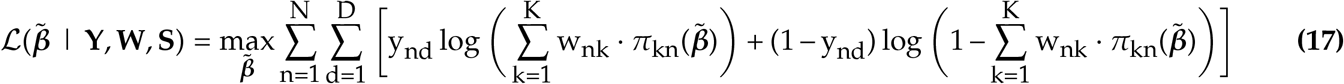

As this log-likelihood expression contains the log of a sum, it is difficult to optimize with respect to 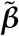. Therefore, we use an Expectation-Maximization to iteratively define lower bound functions that we optimize with respect to 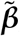, which allows solving (17).

### Expectation-Maximization algorithm to solve CLIFF

By using the concavity of the log through Jensen inequality, we can define a lower bound of (17) (more details on this in the Supplemental Methods) :

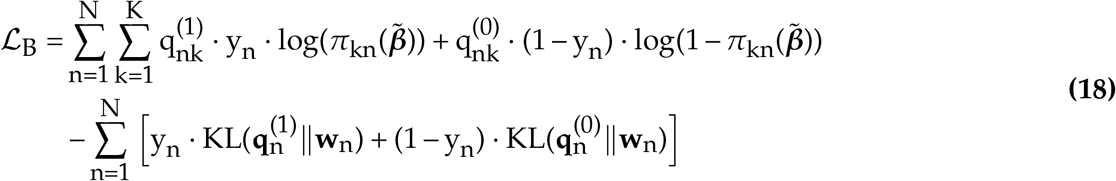

In order to solve (17), we will iterate over the Expectation step or E-step where we update 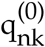 and 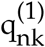 in (18), and the Maximization step or M-step, where we maximize (18) with respect to 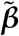. We can therefore define our Maximization algorithm as follow :

#### E-step

During E-step, we define our lower bound function at step t, 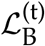 by defining 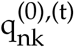,and 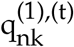 using 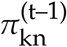 defined at previous step :

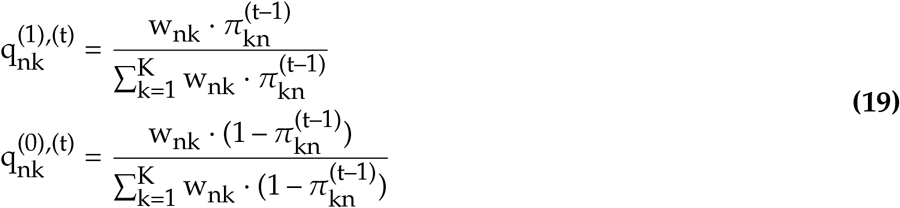

This gives us the lower bound function of (18), tight to the original log-likelihood (17) when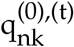 and 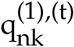 satisfies (19). For a precise description of how 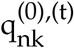, and 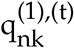, were defined, see Supplemental Methods.

#### M-step

During M-step, we optimize (18) with respect to 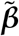, considering the variables **q**^(0),(t)^ and **q**^(0),(t)^ defined during E-step as constants. Also, the two last term of (18) do not depend on 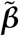, thus we can solve the following problem which will produce the same optimal 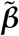 coefficients than if solving (18) :

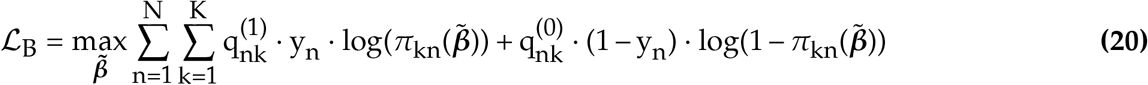

Interestingly, (20) can be solved with standard libraries using a weighted logistic regression model, where we define weights as being 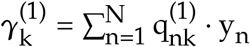 and 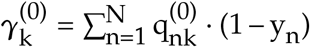.

We optimize 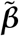 coefficients through a ten-fold cross-validation (taking 90% of the samples N in the training set, and 10% in the testing set), and using early-stopping to terminate the EM algorithm. Optimized 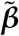 coefficients allow to compute optimized values for *π*_kn_, our estimate of cell-subtype drug sensitivity.

### CLIMB-SC - bulk deconvolution at single-cell level

The bulk deconvolution performed by CLIMB has the particularity of estimating coefficient directly on single-cells from a reference scRNA-seq dataset. Instead of transforming these coefficients into cell-subtype proportions as in Methods (3), we can compute normalized coefficients 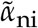 with the following formula :

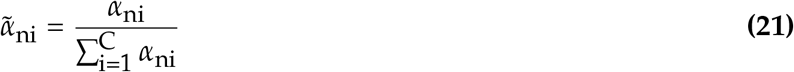

With *α*_ni_ being the coefficients fitted with GLMNET library. We therefore call this *single-cell bulk deconvolution*, which links every bulk RNA-seq n to every single-cells i from a scRNA-seq dataset, through what we will call *single-cell proportions* 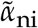. These *single-cell proportions* are thus normalized coefficients fitted by CLIMB, as shown above in (21). Thus we obtain that 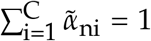 As 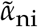 sum up to 1, they can be used by CLIFF as if they were cell-subtype proportions.

### CLIFF-SC for drug sensitivity inference at single-cell level

By slightly modifying CLIFF method and by linking it with the ouput of CLIMB-SC, we propose CLIFF-SC, a method providing drug sensitivity estimates at single-cell resolution. If the input dataset is composed of paired bulk RNA-seq and drug sensitivity screening data, a so-called *bulk deconvolution at single-cell level* is performed using a reference scRNA-seq dataset (cf. CLIMB-SC above).

Thus, as in (15), we model the observed ratio of cells y_nd_ that survives in a mixed population of cells as an average of single-cell survival probabilities *π*_id_, associated with the bulk sample n, weighted by the coefficients 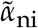 :

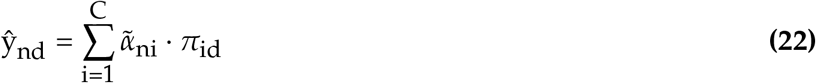

With C being the number of cells contained in a scRNA-seq reference datasets. Thus, we model the single-cell drug sensitivity *π*_id_, which is also modeled with a sigmoid function, similar to (16) :

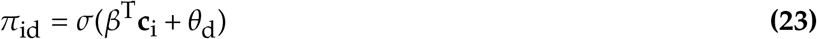

Where **c**_i_ is the ith row of the single-cell expression matrix **C**, thus **c**_i_ is a vector of length G, representing the gene expression of cell i. Similarly as in the cell-subtype-centered model, we can derive a log-likelihood function :

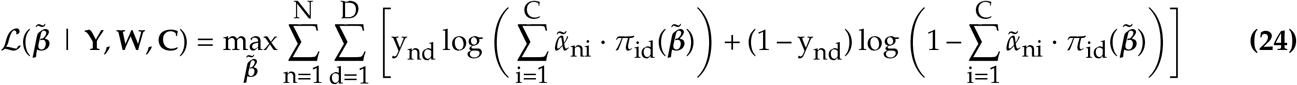

A lower bound can be derived from (24), which gives :

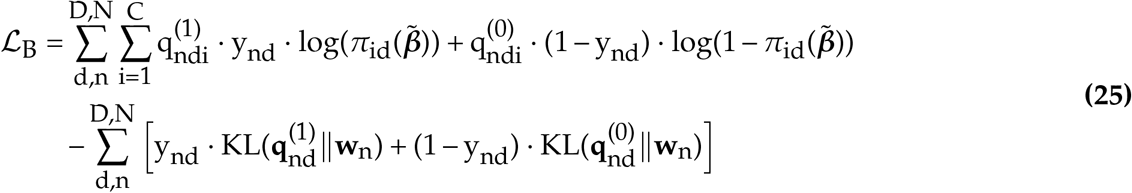

Then, the Expectation-Maximization algorithm can initialized with all coefficients 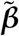 at 0. This implies that 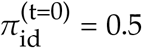 for all i, d. During E-step, we compute values for 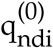 and 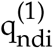 variables :

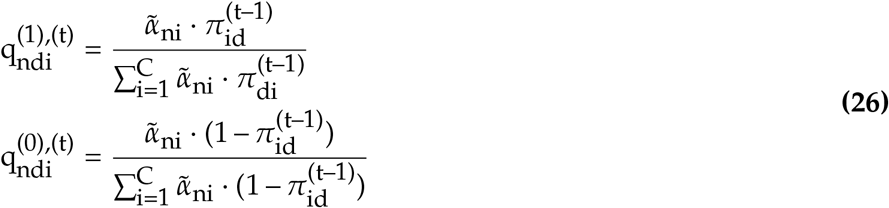

Then, we optimize 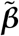 coefficients at each step (t), during the so-called M-step :

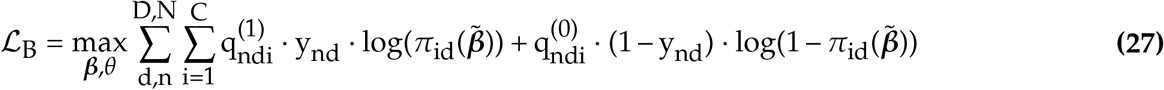

And we run the EM algorithm in a similar way than shown above for CLIFF.

### Simulation of pseudo-bulk samples from scRNA-seq

We generated pseudo-bulk samples by summing up single-cell expression vectors per gene to obtain an artificially generated “pseudo”-bulk RNA-seq. As the precise amount of cells from each cell subtype added to the mix is known, we thus precisely know the cell-subtype proportions of the mixture and can benchmark deconvolution algorithms efficiently. We generated two datasets of pseudo-bulk in this work, both based on single-cells found in the Van Galen et al. dataset.

First, we used all single cells from the same sample (thus, the same patient), which we summed up to obtain a patient-specific pseudo-bulk RNA-seq sample. Thus, we define a variable *ϕ*_ni_ as a binary output variable indicating if a cell i belongs to patient n. Thus, a pseudo-bulk samples is a vector of length G containing the gene expression summed over all cells from patient n :

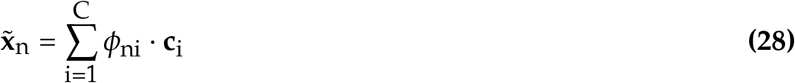

where **c**_i_ is a vector of length G representing the gene expression of cell i. We can therefore easily get the cell-subtype proportion w_nk_, by simply counting the number of cells from each cell subtypes and dividing by the total number of cells in patient n. For this we introduce an indicator vector ***τ***_k_, which is of length C and equal to 1 whenever cell i is of type k.

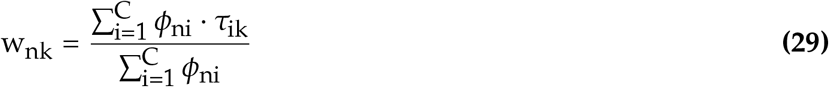

Van Galen et al. dataset offers only a limited number of patients (16 AML patients at day 0 pre-treatment). To simulate an *in-silico* a dataset of similar size to the beatAML cohort, we generated 200 pseudo-bulk samples from the Van Galen et al. dataset allowing a random mix of cells from multiple patients. First, we generated 200 random cell-subtype proportions using a Dirichlet distribution. Thus, we assume that cell-subtype proportions **w**_n_ ∼ Dir(***α***), with ***α*** being a vector of length K, containing one value for each cell subtype k. We fitted ***α*** using cell-subtype proportions in Van Galen et al. dataset with the R library *feralaes/dampack*. Then, we generated cell-subtype proportions for 200 pseudo-bulks, with the R library *DirichletReg* and the ***α*** previously fitted. Cells from Van Galen et al. dataset were selected with probabilities corresponding to cell-subtype proportions. To use a realistic number of cells to generate our pseudo-bulk samples, we used a gamma distribution fitted on Van Galen et al. dataset using *fitdistr* function from R library *MASS*. In the end, we generated a matrix of pseudo-bulk expression 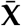 of size (N *×* G) with N = 200, together with associated true cell-subtype proportions 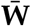of size (N *×* K).

### Simulation of drug sensitivity screening data

To simulate drug screening data, we assumed that the drug sensitivity observed on a mix of cells corresponds to the sum of the cell-subtype drug sensitivity constituting its bulk weighted by their respective proportions. Additionally, we added Gaussian noise with a signal-to-noise ratio of 100. Thus, the simulated survival rate for pseudo-bulk n, in cell-subtype k exposed to drug dose d was generated with the following formula, given the proportions :

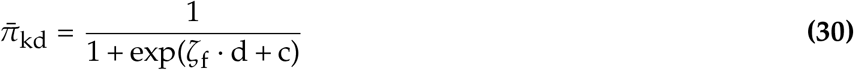

with *ζ*_f_ being a randomly generated number, with *ζ*_f_ ∈ [1, 2] if f = 1 (low sensibility cell-subtypes - resisting cells) and *ζ*_f_ ∈ [5, 6] if f = 2 (high sensibility cell-subtypes). Here we did not use the drug dose d, only the integer number related to drug dose d = {0, 1, …, 6}, as we considered seven drug doses. We arbitrarily defined three cell subtypes with high sensitivity and the rest having low sensitivity. The constant c added was set to c = –8 as it provided visually convincing results. Thus 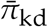 follows a sigmoid function that depends on drug dose d but in a way that is slightly different than in our model. Then, we generated the survival rate at the bulk level using the same formula as in our model but adding gaussian noise to it :

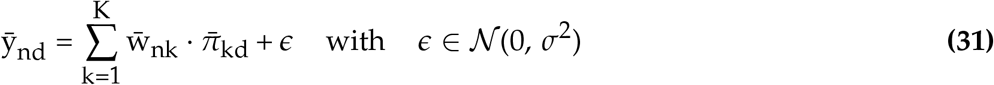

and *σ* was chosen to generate a signal-to-noise ratio arbitrarily chosen at 100.

### Area under the curve for summarizing drug sensitivity of multiple data points

As CLIFF produces D predictions for each cell subtype, and CLIFF-SC produces D predictions for every single cell, it is desired to summarize these D data points into a single value. The goal is to summarize cell-subtype or single-cell drug sensitivity. To summarize drug sensitivity from D different drug doses, we normalize drug doses so that the minimum dose = 0 and the maximum dose = 1. As the cell survival rate is ∈ [0, 1], we can compute an area under the curve (AUC) which will be ∈ [0, 1]. Conversely, we get an AUC of 1.0 if all cell survives and an AUC of 0.0 if all cells die, taking every dose into account. We computed the AUC using the function *computeAUC* from the R package *PharmacoGx*.

### Culture conditions of cancer cell lines

Human cancer cell lines (K562, HL60, THP1, SUDHL4) were purchased from the American Type Culture Collection (ATCC; Rockville, MD, USA) and cultured at 37°C in a humidified 5% CO2 environment in RPMI 1640 (HCC38) supplemented with 2mM L-glutamine, 2 g/L D-glucose, and 10% fetal bovine serum (FBS; Gibco ThermoFisher).

### Pharmaceutical compounds

Stock solutions for every agent (Table2) were prepared using DMSO (Sigma-Aldrich; stored at 80°C), further diluted in H2O to the appropriate concentration, plated in Greiner 96-well flat bottom (REF) and the plates were stored at 20°C. The pharmaceutical compounds were screened at eight concentrations (Log concentration M = -9.205031395, -8.699881554, -8.130127907, -7.70415926, -7.209309101, -6.70415926, -6.125806332, -5.699837685, -5.204987526, -4.699837685) using a Log2-fold dilution series with matched DMSO concentration vehicle controls.

**Table 1.** Table of eight coumpounds used in the *in-vitro* drug sensitivity experiment.

| CatalogID | Name | Supplier |
| --- | --- | --- |
| A3656 | 5-Aza-2'-deoxycytidine | Sigma |
| C3350000 | Cytarabine | Sigma |
| CDS022173 | Imatinib | Sigma |
| CDS023093 | Nilotinib | Sigma |
| HY-15142 | Doxorubicin (hydrochloride) | MedChemExpress |
| O9512 | Oxaliplatin | Sigma |
| R0395 | Rapamycin | Sigma |
| Y0000378 | Mitomycin | Sigma |

**Table 2.** Table of cell line proportions added in tubes manually to generate the mixtures of cell lines.

| Cell line | Mixture1 | Mixture2 | Mixture3 | Mixture4 | Mixture5 | Mixture6 |
| --- | --- | --- | --- | --- | --- | --- |
| SUDHL4 | 0.33 | 0.07 | 0.00 | 0.61 | 0.188 | 0.20 |
| K562 | 0.09 | 0.11 | 0.222 | 0.04 | 0.317 | 0.26 |
| HL60 | 0.26 | 0.57 | 0.333 | 0.05 | 0.0198 | 0.12 |
| THP1 | 0.32 | 0.25 | 0.444 | 0.30 | 0.475 | 0.42 |

### Bulk proportion for bulk RNAseq and drug sensitivity

Six bulks were created by mixing the four cell lines with the following proportions. Pure and mixed cell populations were subjected to RNA extraction and drug sensitivity immediately after mixing.

### Cell viability assay

Cells were plated in the 96-well plates containing chemical compounds previously equilibrated at 37C, at a density of 1×104 cells per well in 100µl culture medium. Cell viability was assessed after drug treatment for 24, 48, 72 or 96 in duplicates hours by incubating the cells with PrestoBlue™ Cell Viability Reagent, (Thermo Fisher Scientific - Life Technologies A13261). PrestoBlue was at room temperature before adding to the cells in each well with. The cells were incubated with 10µl PrestoBlue (10% of cell culture volume) for two hours at 37°C. The fluorescence was measured by the bottom with a 560nm excitation filter and a 590nm emission filter in a BioTek Neo HTS microplate. The acquisitions were performed with the following parameters: Gain: 54, Light Source Xenon Flash, Lamp Energy: High, Read Speed: Normal, Delay 0 msec, Mesurements/Data Point: 10, Read Height 5.25mm.

### RNA extraction and sequencing

RNA-seq Total RNA from cell lines was isolated with NucleoSpin™ RNA Plus kit (Machery-Nagel). cDNA was prepared with Maxima Reverse Transcriptase (Thermo Scientific). Sequencing libraries were performed with Illumina Truseq Stranded mRNA LT kit. Reads were mapped to the human (hg19) genome using Hisat2. Counts on genes were generated using featureCounts. Normalization for sequencing depth has been done using the counts on genes as library size using the TMM method as implemented in the limma package of Bioconductor.

### Single Cell RNAseq (MULTIome-Seq)

Single cell RNAseq was performed using 10x Genomics Chromium Single Cell Controller following the 10x Genomics Multiome (scRNA/scATAC-seq) demonstrated protocol: Chromium Next GEM Single Cell Multiome ATAC + Gene Expression User Guide (CG000338). To note, experimental parameters specific to the experiment were 3,000 for the Targeted Nuclei Recovery and 5 minutes for the lysis time.

### Single-cell RNA-seq analysis

Cellranger-arc was used to obtain counts on genes using default parameters. Counts were obtained using cellranger using hg19. Only uniquely mapped reads on genes that were expressed in at least 1% of the samples were kept. Seurat’s SCTransform was used to normalize the data and correct for mitochondrial percentage and total number of reads biases.

## DATA AVAILABILITY

The data produced for the in-vitro experiment shown in Fig. 2 and Fig. 4 can be found on a Gene Expression Omnibus repository (GEO): GSE217924 (superserie). The bulk RNA-seq of the mixtures is available on the GSE217919 repository and the single-cell RNA-seq data on the GSE217922 repository. The rest of the study was relying on re-analysis of existing data available at the following GEO locations: Van Galen et al. (GSE116256), Naldini et al. (GSE185834), Lee et al. (GSE144735), Khaliq et al. (GSE200997), Gray et al. (GSE180878), Wu et al. (GSE176078), Jerby et al. (GSE115978), Tirosh et al. (GSE115978), Neftel et al. (SmartSeq2 and 10X data in GSE131928). BeatAML dataset is available via dbGaP study ID 30641 and accession ID phs001657.v1.p1.

## CODE AVAILABILITY

The code is freely accessible on GitHub. CLIMB and CLIFF code can be accessed via an R library, ‘ClimbTheCliff’, available via the devtools command devtools::install_github(‘alexdray86/ClimbTheCliff’). Documentation are available on https://alexdray86.github.io/CLIMB/build/index.html and https://alexdray86.github.io/CLIFF/build/index.html website.

## ACKNOWLEDGMENTS

This work was supported by grants from the Swiss National Science Foundation and the European Research Council (KRABnKAP, no. 268721; Transpos-X, no. 694658), the Personalized Health and Related Technologies (PHRT) and Swiss Data Science Center (SDSC) strategic focus areas of the Swiss Federal Institutes of Technology, the Swiss Personalized Health Network (SPHN) initiative of the Swiss Academy of Medical Sciences (project no. 2017-407), The Swiss Cancer League and the Aclon Foundation to D.T. We also extend our gratitude to Daria Korotkova, Cyril Pulver, and Paola Malsot, for their valuable scientific discussions. Additionally, I acknowledge my wonderful son, Matisse Coudray, for the inspiration he has provided.

---

**Supplementary Table 1.**
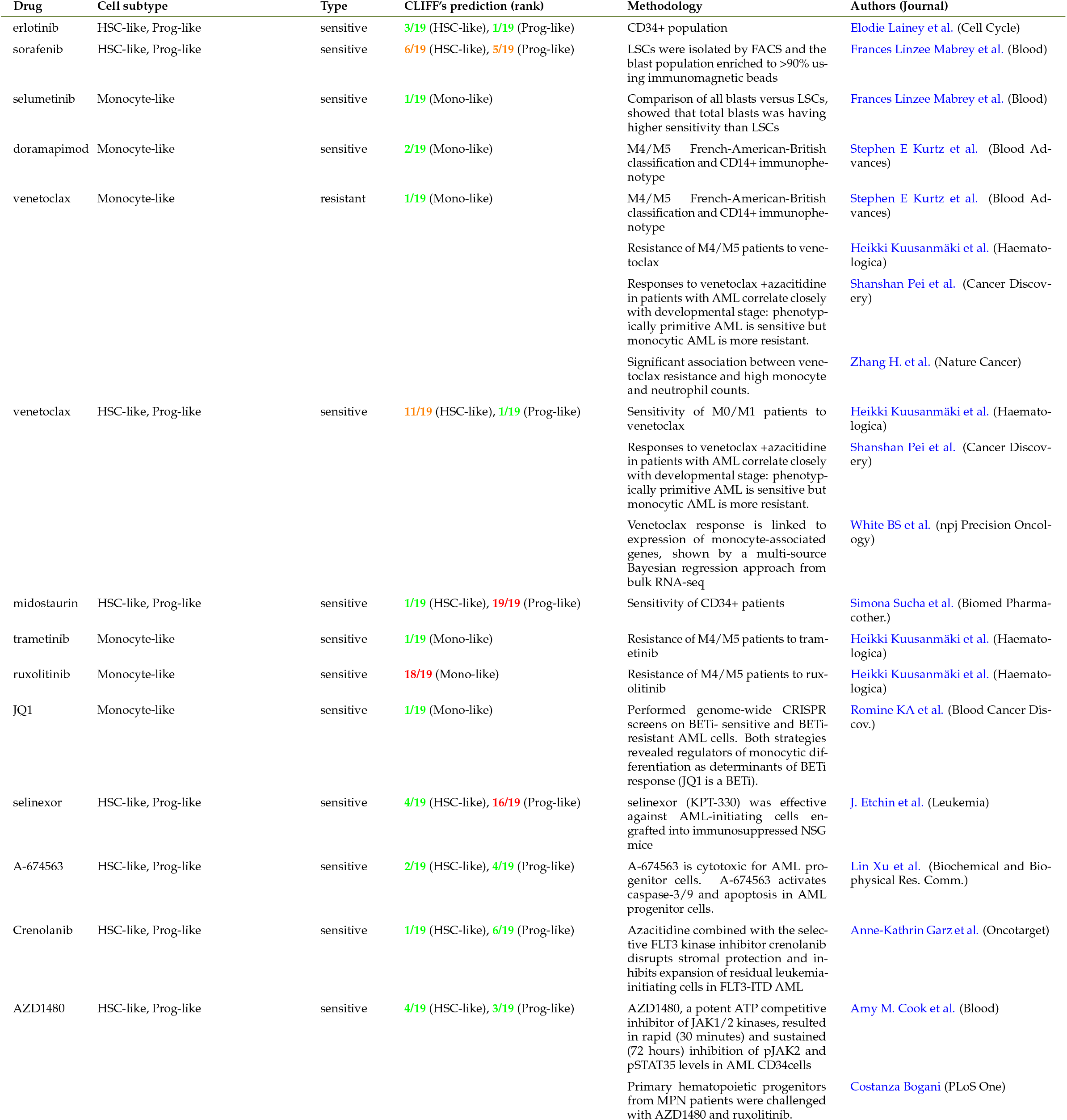
Documented cell-subtype drug sensitivity in Acute Myeloid Leukemia. Green signifies CLIFF’s predictions aligned with the literature (25% most sensitive/resistant). Red denotes CLIFF’s predictions conflicting with the literature (25% less sensitive/resistant). Orange represents uncertain connections, with values predicted between the 25% and 75% quartiles.

**Supplementary Figure 1.**
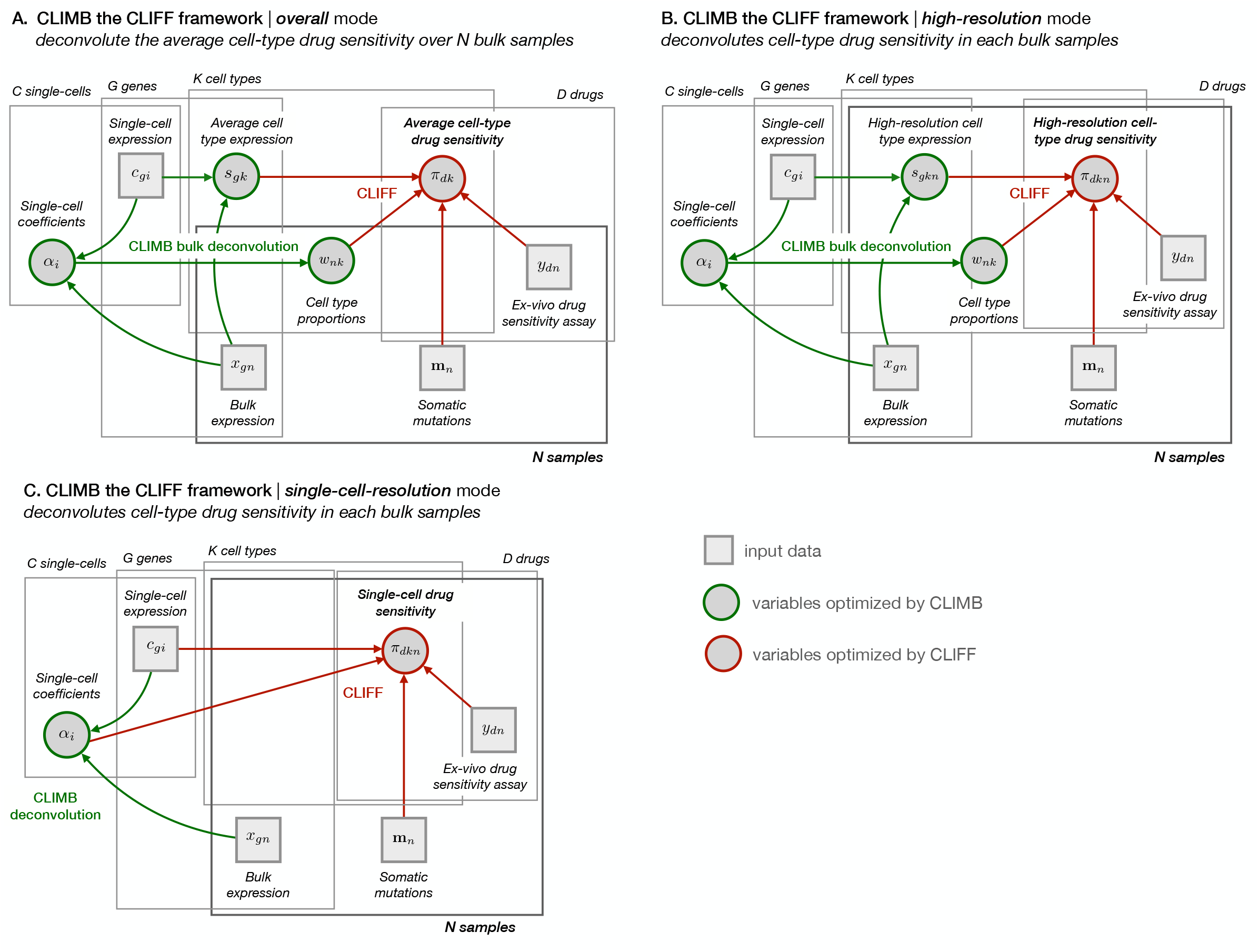
(**A**) Diagram of CLIMB / CLIFF input / output variables in overall mode. (**B**) Same as (**A**) in high-resolution mode: cell-subtype expression and cell-subtype drug sensitivity is predicted in each bulk sample.

**Supplementary Figure 2.**
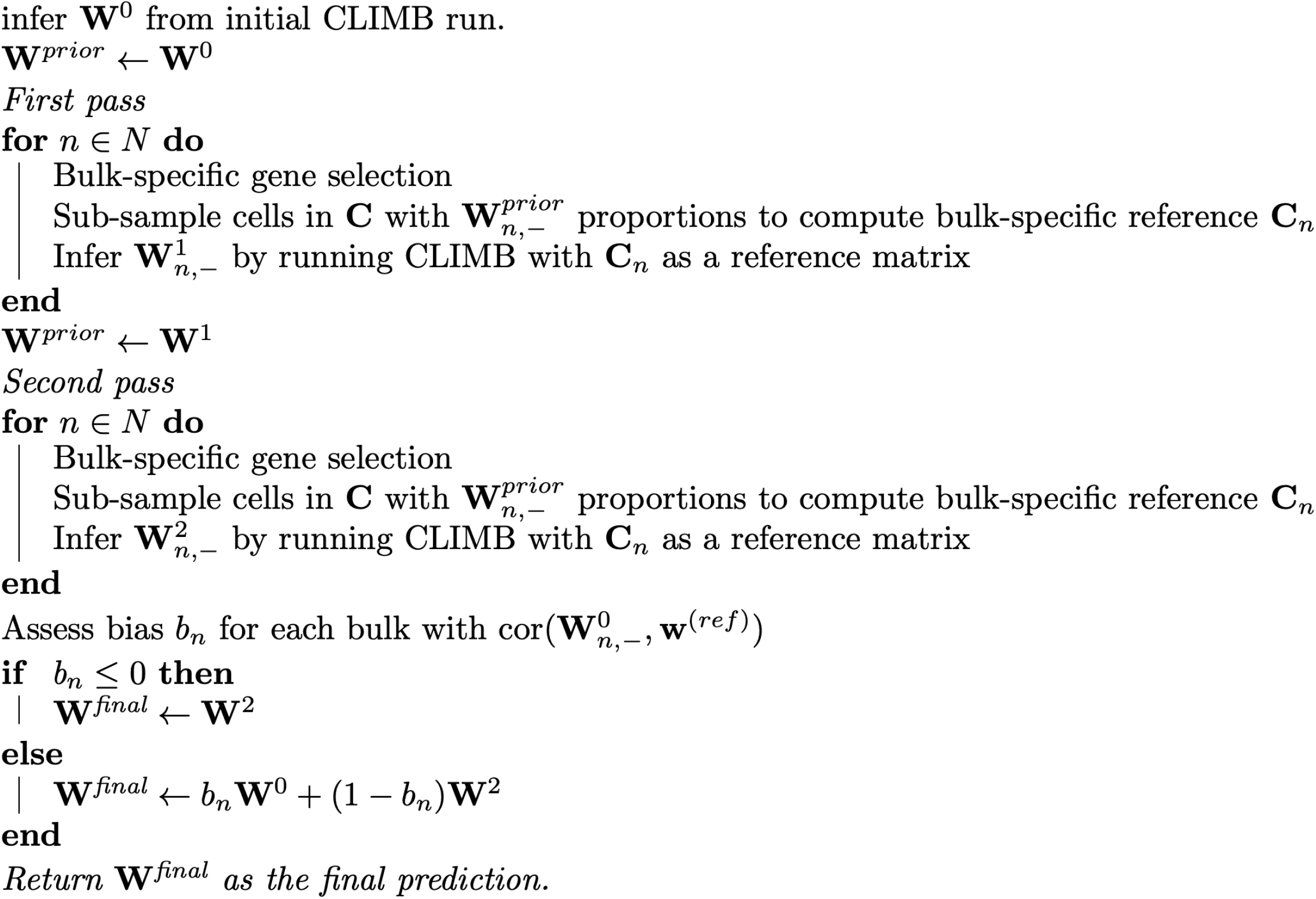
CLIMB 2-pass algorithm. The algorithm subset cells in the reference single-cell matrix C two times. W represents the predicted cell-type proportions, of size NxK, N being the number of bulk sample and K the number of cell types. The first pass allows to subsample cells in the reference matrix, which is then used during the second pass to infer the cell subtype proportions. This process is iterated twice.

**Supplementary Figure 3.**
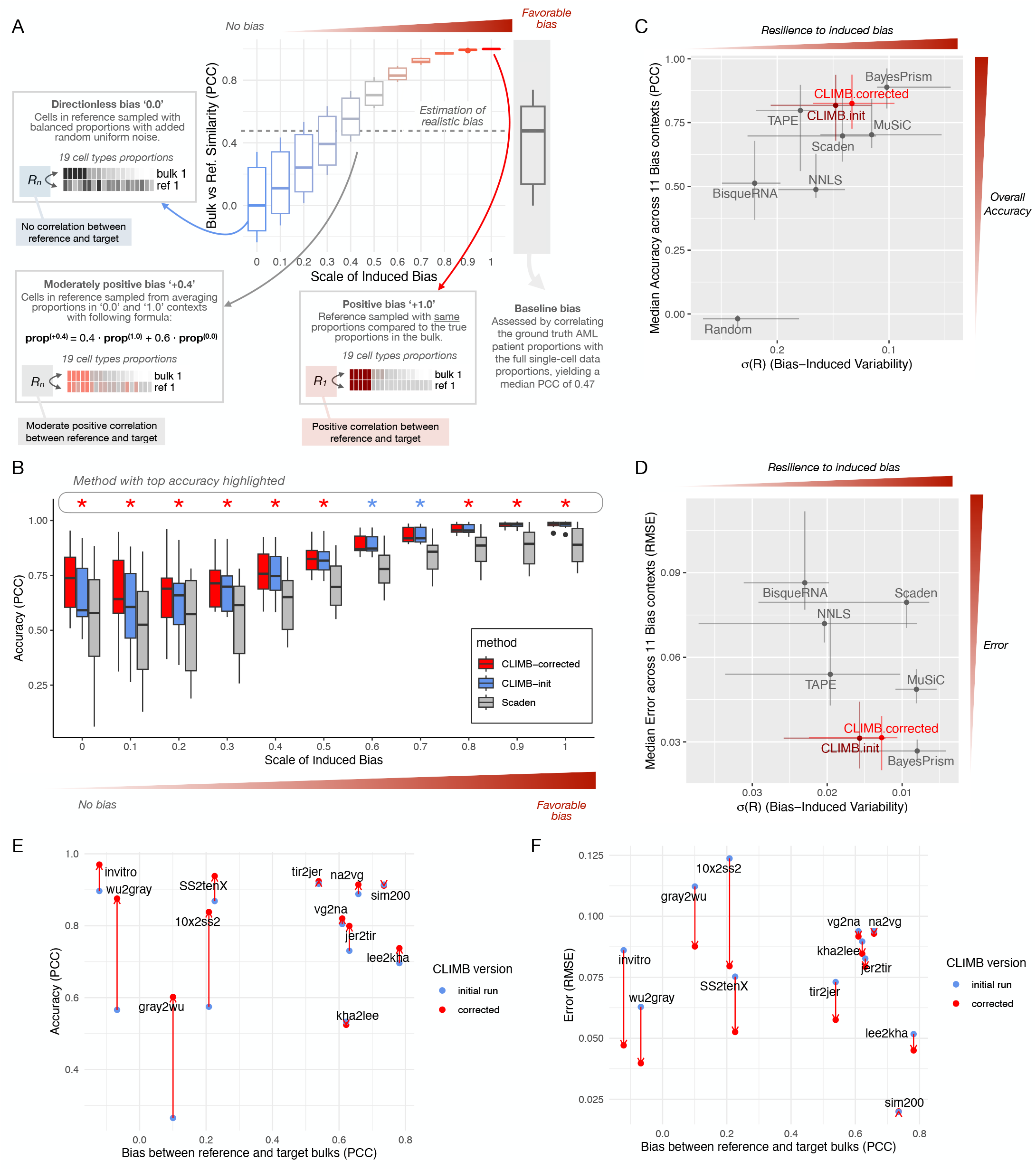
(**A**) Illustration of the simulated pseudo-bulks with varying biases between reference and target proportions. On the right, we show the bias by comparing Van Galen et al. data with its own pseudo-bulks. (**B**) Accuracy, shown as a correlation coefficient (PCC), comparing true vs. predicted cell type fractions across 11 simulated scenarios for CLIMB-init, CLIMB-corrected, and Scaden methods. (**C**) Comparative accuracy and resilience to bias for all deconvolution methods. Resilience to bias is assumed to be equal to the variability in accuracy across different biases. (**D**) Same as (C), but showing the results as an error (RMSE). (**E**) Comparison between CLIMB-init and CLIMB-corrected deconvolution accuracy (PCC) across all analyses (in vitro and pseudo-bulks). (**F**) Same as (E), but showing the error instead.

**Supplementary Figure 4.**
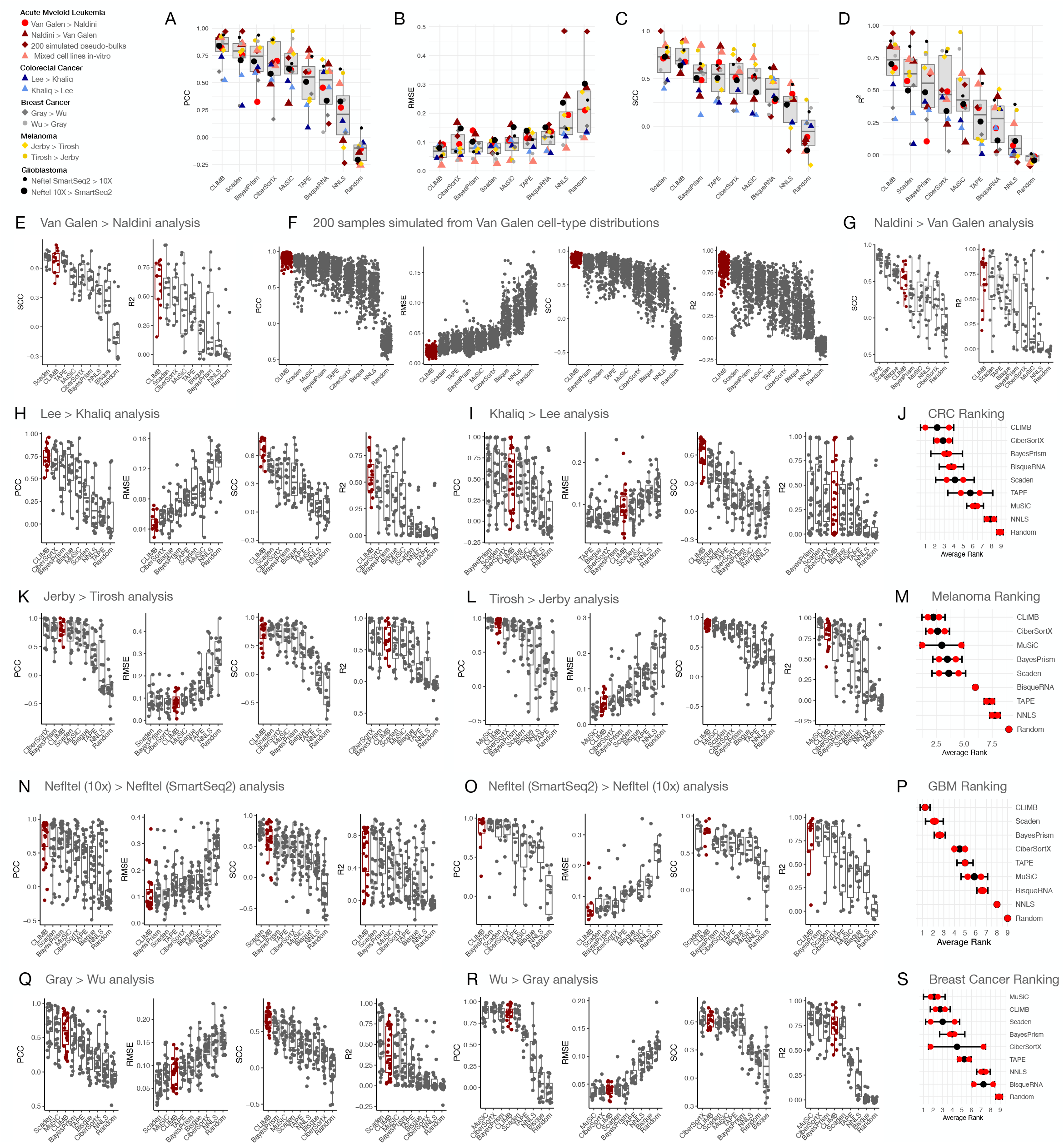
(**A-D**) Combined pseudo-bulk deconvolution analysis across datasets showing four accuracy metrics. (**E**) Pseudo-bulk deconvolution analysis across datasets (Van Galen > Naldini) displaying accuracy as Spearman correlation coefficient / R-square value per sample (one data point per sample), comparing true and predicted cell subtype proportions. (**F**) Deconvolution method prediction accuracy for 200 simulated pseudo-bulk samples derived from the Van Galen et al. dataset. Results presented as boxplots, per sample level, for PCC, SCC, RMSE, and R-square metrics. (**G**) Similar to (**E**), but for Naldini > Van Galen cross-dataset deconvolution analysis. (**H-J**) Cross-dataset deconvolution analysis in colorectal cancer, depicted as boxplots for PCC, SCC, RMSE, and R-square metrics, covering Lee > Khaliq analysis (**H**) and Khaliq > Lee analysis (**I**). (**J**) Average rank over 4 metrics considering Lee > Khaliq and Khaliq > Lee cross-dataset analyses. (**K-M**) Melanoma cross-dataset deconvolution analysis, showcasing the same metrics as (**H-J**), for Jerby > Tirosh analysis (**K**) and Tirosh > Jerby analysis (**L**). (**N-P**) Glioblastoma cross-dataset deconvolution analysis, for Neftel (10X) > Neftel (SmartSeq2) analysis (**N**) and Neftel (SmartSeq2) > Neftel (10X) analysis (**O**). (**Q-S**) Cross-dataset deconvolution analysis in breast cancer, for Gray > Wu analysis (**Q**) and Wu > Gray analysis (**R**).

**Supplementary Figure 5.**
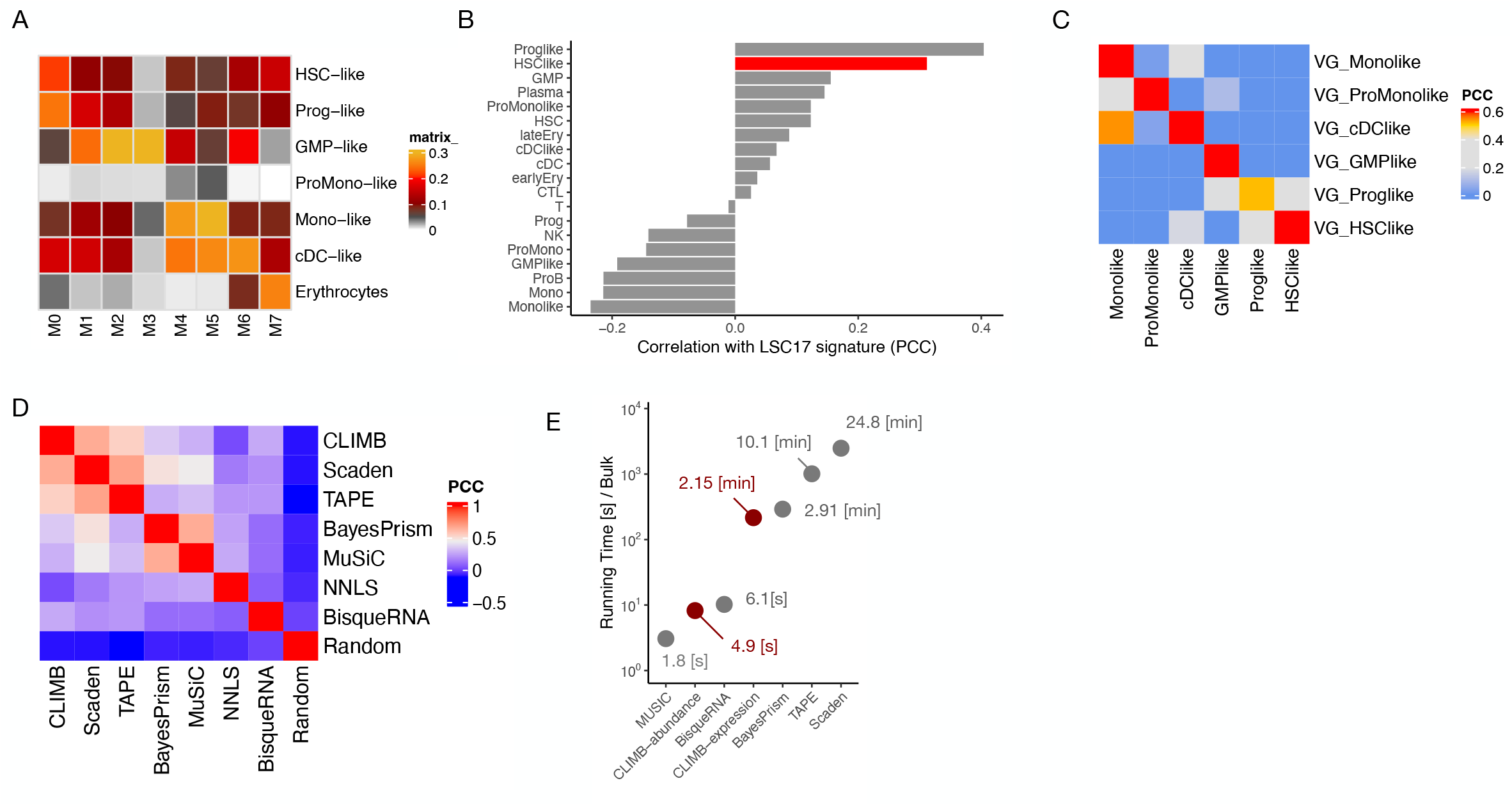
(**A**) Average cell type proportions in different categories of FAB patients. (**B**) Correlation (PCC) between CLIMB-predicted cell subtype proportions and LSC17 signature. (**C**) Correlation (PCC) between Van Galen cell type signature and cell type proportions estimated by CLIMB. (**D**) Correlation (PCC) between cell subtype abundance estimated by different deconvolution methods in merged BeatAML, Leucegene, TCGA-AML datasets. (**E**) Running time of various deconvolution methods.

**Supplementary Figure 6.**
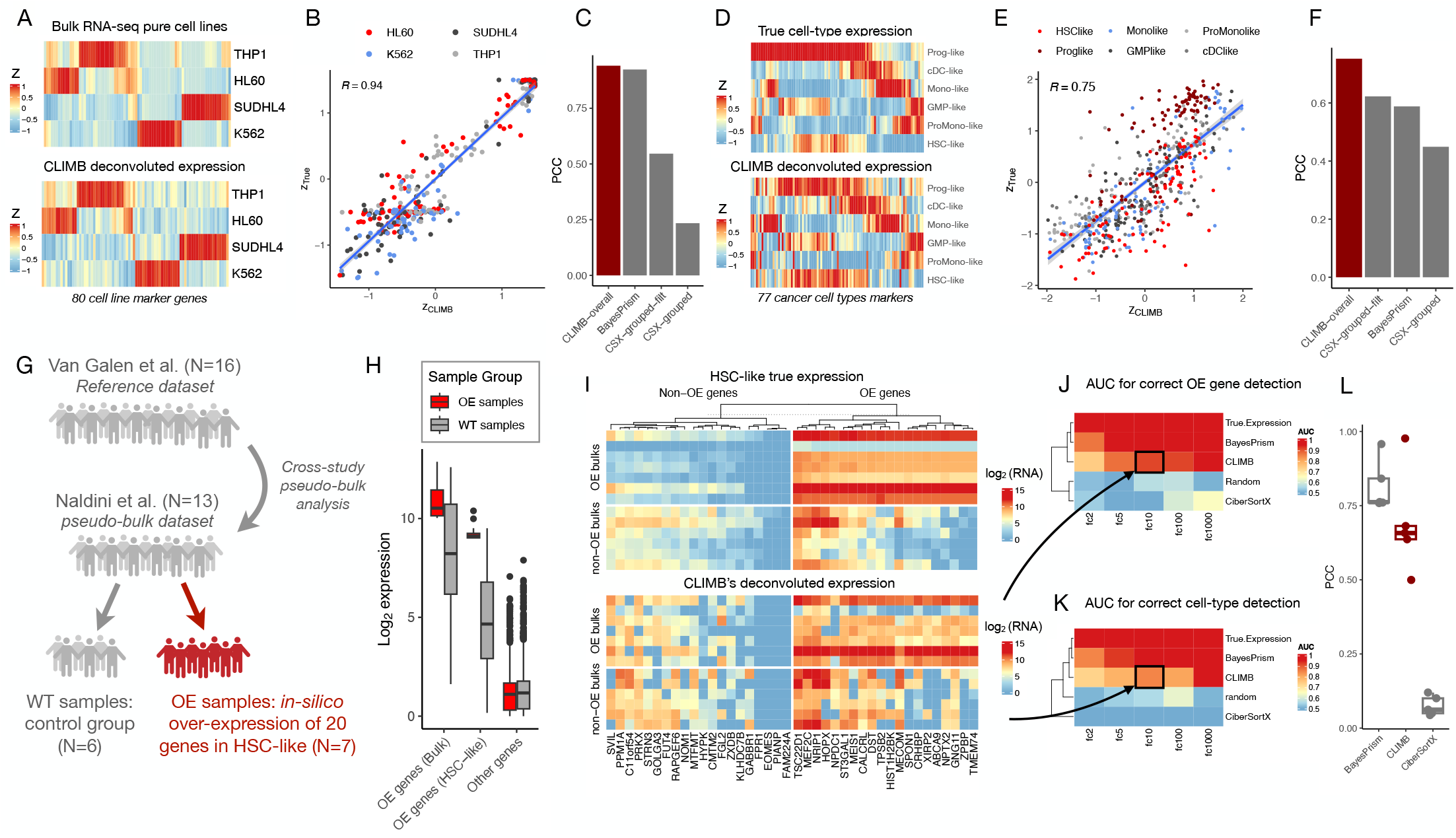
Deconvolution of cell subtype Expression and Simulated Over-Expression Evaluation. (**A**) Heatmap comparing standardized true expression from bulk RNA-seq on pure cell lines (top) with deconvoluted expression from CLIMB based on bulk RNA-seq of cell line mixes (bottom). (**B**) Scatter plot comparing standardized true expression (y-axis) and CLIMB’s deconvoluted expression (x-axis) from bulk RNA-seq. (**C**) Accuracy as PCC comparing true expression from bulk RNA-seq and deconvoluted expression from CLIMB, BayesPrism, CiberSortX (group mode, filtered and unfiltered). (**D**) Heatmap showing standardized true expression from pseudo-bulks (top) and CLIMB’s deconvoluted expression of pseudo-bulks (bottom). (**E**) Scatter plot of deconvoluted expression (x-axis) versus true expression (y-axis) for pseudo-bulks. (**F**) Accuracy as PCC comparing true expression from bulk RNA-seq and deconvoluted expression from CLIMB, BayesPrism, CiberSortX (group mode, filtered and unfiltered). (**G**) Schematic of cross-dataset analysis with over-expression of 20 genes in 7/13 bulk samples and in HSC-like cell subtype. (**H**) Fold changes observed at bulk and cell-subtype levels when inducing single cells with a fold change of 10 before pseudo-bulk generation. Control genes are also compared. (**I**) Comparison between true and CLIMB’s deconvoluted expression for HSC-like with over-expressed genes. Top heatmap displays true expression, with 20 over-expressed genes (right) and 20 random non-over-expressed genes (left). Overexpression samples are above, WT samples below. Bottom heatmap shows CLIMB’s deconvoluted expression with the same gene and sample order. (**J**) Accuracy to detect correct over-expressed genes in HSC-like, using*·* –log_10_(padj) · FC by DESeq2, presented as an average AUC-ROC over 20 genes. (**K**) Similar to (**J**), but showing accuracy to detect the correct cell subtype among cancer cell subtypes for each over-expressed gene, averaged over 20 genes. (**L**) Accuracy as PCC comparing true expression with CLIMB, BayesPrism, and CiberSortX deconvoluted expression.

**Supplementary Figure 7.**
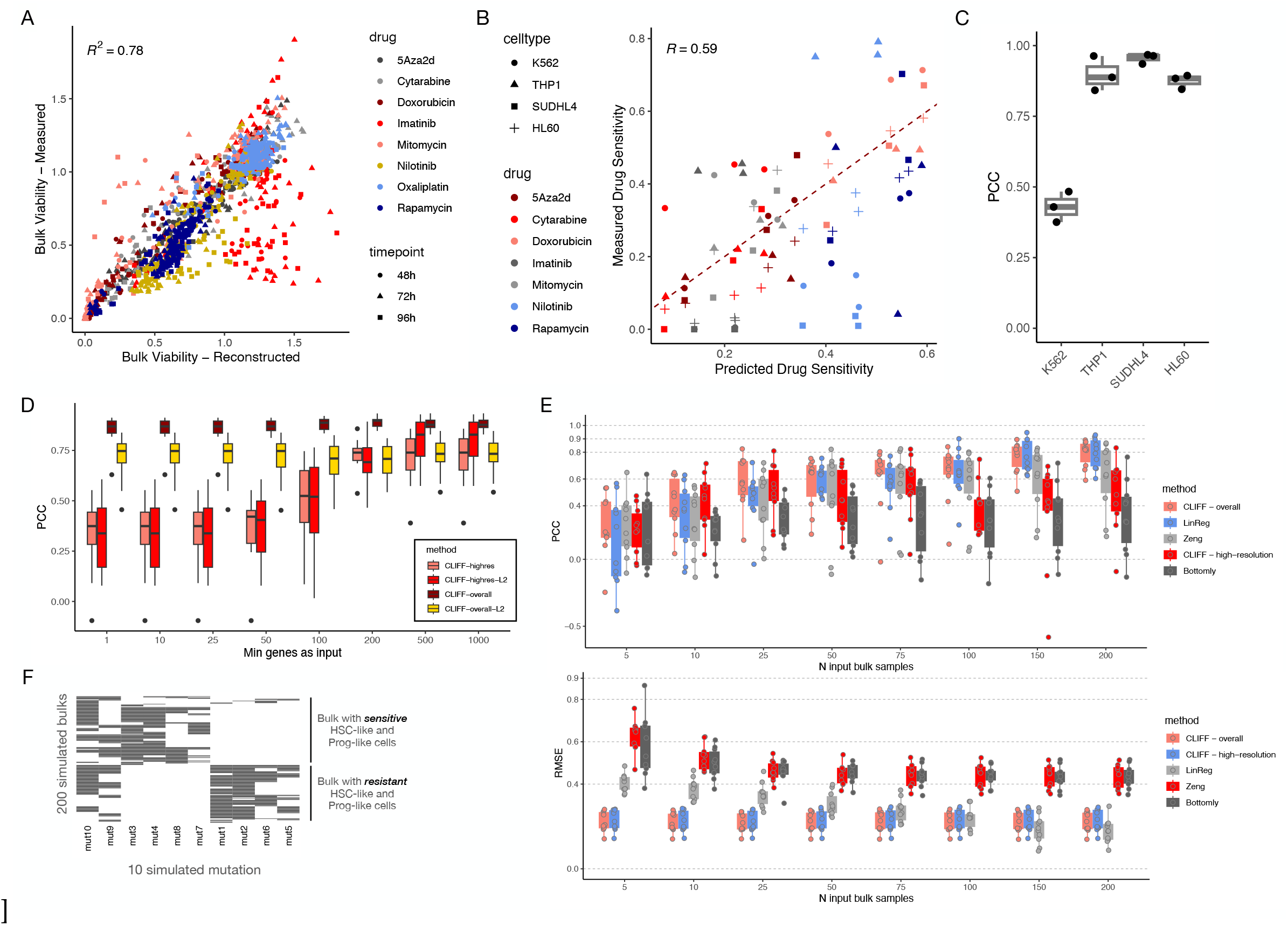
(**A**) Comparative analysis of true bulk viability (measured for 3 timepoints, 7 drugs, and 10 doses) against reconstructed bulk viability, using experimentally-assessed viability of pure cell lines comprising the bulks. (**B**) True cell line drug sensitivity for multiple timepoints, cell lines, and drugs, juxtaposed with CLIMB’s high-resolution mode predictions (subsequently averaged at cell type level). (**C**) CLIFF’s predictive accuracy presented as Pearson correlation coefficient (PCC), grouped by cell lines. (**D**) Accuracy comparison of CLIFF with varying numbers of input genes, based on the 200 simulated bulks. We evaluated four modes for CLIFF: CLIFF-overall, CLIFF-overall-L2 (CLIFF-overall with additional L2 regularization), CLIFF-highres (subsequent average at cell subtype level), and CLIFF-highres-L2 (CLIFF-highres with L2 regularization). (**E**) Predictive accuracy depicted as PCC for deconvolution of cell subtype proportions in 200 simulated pseudo-bulks, featuring CLIFF-overall, CLIFF-highres, Karakalsar’s approach (linear regression using MuSiC-derived proportions as input matrix), and Bottomly’s approach (correlation between cell-subtype score obtained from PC1 of cell subtype signature, and bulk drug sensitivity). (**F**) Simulated mutational landscape for 200 simulated bulks employed in high-resolution drug sensitivity deconvolution.

**Supplementary Figure 8.**
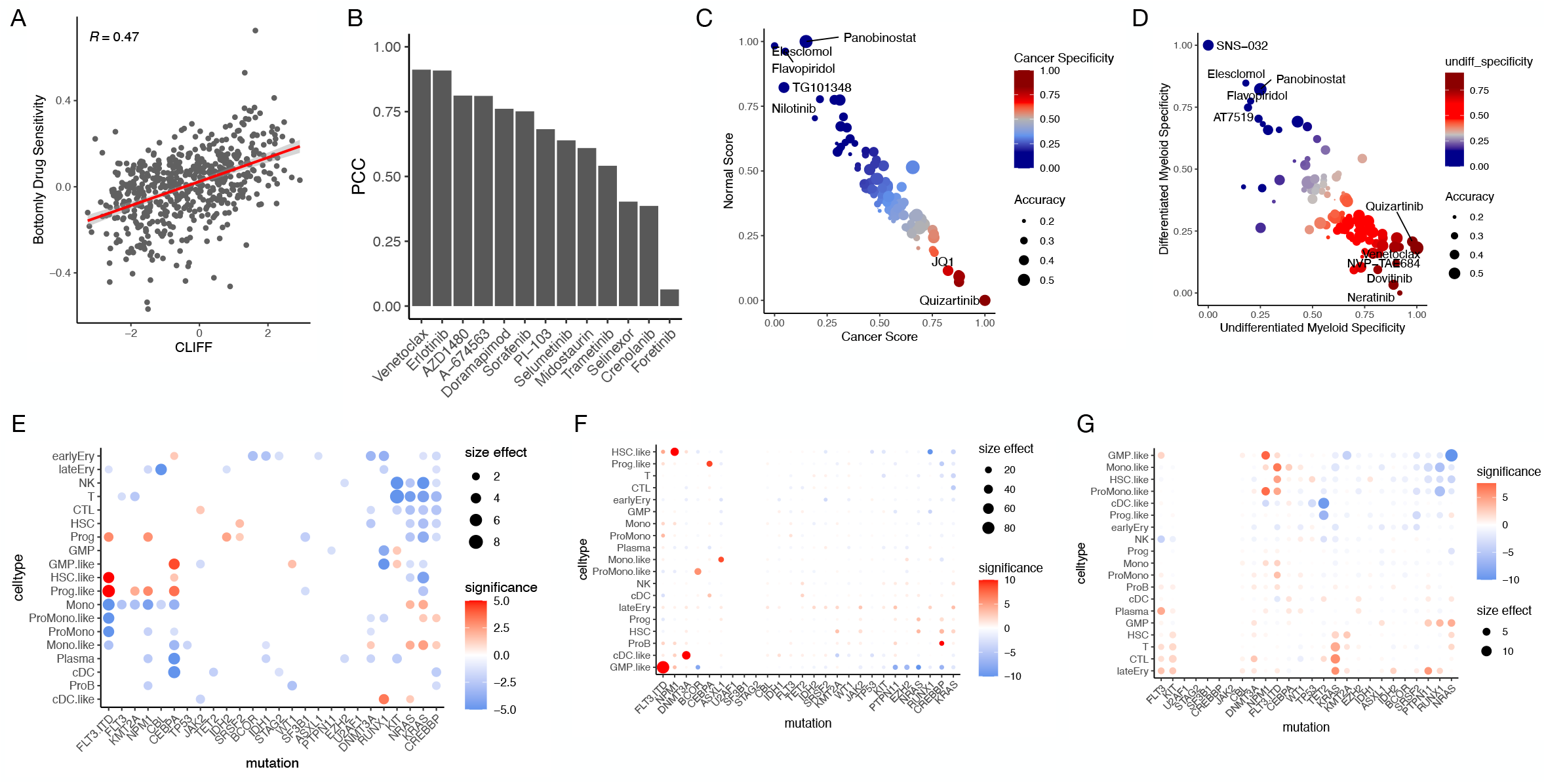
(**A**) Scatter plot illustrating the Pearson correlation coefficient (PCC) between Bottomly’s drug sensitivity predictions (for the 6 cancer cell subtypes defined by Van Galen et al.) and standardized CLIFF-overall predicted AUC (relative values of cell subtype drug sensitivity). Each data point represents a specific cell subtype-drug combination. (**B**) Corresponding to (**A**), this plot displays the overall PCC (averaging 6 data points for the 6 cancer cell subtypes) for the 14 drugs listed in Supplementary Table 1. (**C**) Predicted cancer cell specificity for drugs within the beatAML cohort, as projected by CLIFF-overall. (**D**) Forecasted drug specificity for undifferentiated versus differentiated blasts in the beatAML cohort, according to CLIFF-overall. (**E**) Overall correlations between mutations and cell subtype proportions using CLIMB’s deconvoluted proportions on the beatAML cohort. A set of mutations with at least 5 positive samples is analyzed, with size-effect/significance depicted through t-test-derived measures. Positive correlations are represented in red, negative in blue. The size indicates the size effect (mean proportion difference). (**F**) Correlation between high-resolution deconvoluted drug sensitivity for each cell subtype and mutations in the beatAML cohort, focusing on the A-674562 compound. (**G**) Same as (**F**), but for the venetoclax drug.

